# A technology-agnostic long-read analysis pipeline for transcriptome discovery and quantification

**DOI:** 10.1101/672931

**Authors:** Dana Wyman, Gabriela Balderrama-Gutierrez, Fairlie Reese, Shan Jiang, Sorena Rahmanian, Stefania Forner, Dina Matheos, Weihua Zeng, Brian Williams, Diane Trout, Whitney England, Shu-Hui Chu, Robert C. Spitale, Andrea J. Tenner, Barbara J. Wold, Ali Mortazavi

**Affiliations:** University of California, Irvine, Department of Developmental and Cell Biology, Irvine, CA 92697, USA; University of California, Irvine, Center for Complex Biological Systems, Irvine, CA 92697, USA; California Institute of Technology, Division of Biology, Pasadena, CA 91125, USA; University of California, Irvine, Department of Pharmaceutical Sciences, Irvine, CA 92697, USA; University of California, Irvine, Department of Molecular Biology and Biochemistry, Irvine, CA 92697, USA; Institute for Memory Impairments and Neurological Disorders, University of California, Irvine, CA 92697, USA; Department of Neurobiology and Behavior, University of California, Irvine, Irvine, CA 92697, USA

## Abstract

Alternative splicing is widely acknowledged to be a crucial regulator of gene expression and is a key contributor to both normal developmental processes and disease states. While cost-effective and accurate for quantification, short-read RNA-seq lacks the ability to resolve full-length transcript isoforms despite increasingly sophisticated computational methods. Long-read sequencing platforms such as Pacific Biosciences (PacBio) and Oxford Nanopore (ONT) bypass the transcript reconstruction challenges of short reads. Here we introduce TALON, the ENCODE4 pipeline for platform-independent analysis of long-read transcriptomes. We apply TALON to the GM12878 cell line and show that while both PacBio and ONT technologies perform well at full-transcript discovery and quantification, each displayed distinct technical artifacts. We further apply TALON to mouse hippocampus and cortex transcriptomes and find that 422 genes found in these regions have more reads associated with novel isoforms than with annotated ones. We demonstrate that TALON is a capable of tracking both known and novel transcript models as well as their expression levels across datasets for both simple studies and in larger projects. These properties will enable TALON users to move beyond the limitations of short-read data to perform isoform discovery and quantification in a uniform manner on existing and future long-read platforms.

## INTRODUCTION

Differences in gene expression play a large role role in shaping cell phenotypes and interactions, both during development and in later life. While humans have around 20,000 protein coding genes, they produce at least 100,000 splice isoforms through alternative splicing, and potentially many more^1^. Alternative splicing controls which exons are included in the mature mRNA, thus expanding the number of possible transcripts that a single gene can encode. Some isoforms have vastly different functions and may be highly specific to a particular tissue or temporal stage^2–4^. For instance, alternative splicing of the transcription factor *erbAα* in rats gives rise to one isoform which acts as a transcriptional activator, while a second isoform acts as a repressor^5^. This is a specific instance of an evolutionary strategy whose extent is not yet known, in which differential RNA splicing creates one or more “dominant negative” protein isoforms. Differential RNA isoforms are also important in disease. The *Mapt* gene has isoforms that are known to be differentially expressed in various human neural lineages, and their relative proportions change during progression of Alzheimer’s disease, ultimately leading to the formation of the tangles that kill neurons^6^.

In the best understood cases, alternative splicing is tightly regulated, relying on highly conserved sequence and structure motifs and complex networks of RNA binding protein interactions to define functional isoforms^7^. Disruptions to the splicing process frequently lead to disease, whether in the form of genetic mutations that directly affect splice sites or splicing factors, or more subtle changes that alter the balance between different isoforms^6,7^. As a result, alternative splicing and exon usage in RNA transcripts have long been the subject of great interest in the context of development and disease. In early studies, the preferred methods for characterizing and measuring isoforms were RT-PCR, Sanger sequencing of expressed sequence tags (ESTs), and isoform-specific microarrays^8^. This changed dramatically with the availability of next-generation short-read RNA sequencing, which allows gene expression to be profiled quantitatively in a high-throughput manner^9^. This led to the generation of large reference transcriptome databases for human and mouse cell types and tissues, begininning with ENCODE and rapidly expanding to GTEx and FANTOM^10–12^. In the cancer community, the Cancer Genome Atlas (TCGA) serves as a massive source of RNA-seq data from patient samples^13^.

With the widespread availability of RNA-seq, many efforts have been made to infer isoform usage from short-read data^14^. However, this is intrinsically challenging, as short-read protocols require cDNA transcripts to be sheared into 50-300 bp pieces prior to sequencing. These pieces are far smaller than typical mammalian transcripts, which can be multiple kilobases in length^15^. This means that it is not possible to know the exact combination of exons originally present in each transcript molecule. To get around this, computational methods were developed to reconstruct the transcript models present in a sample and to quantify their abundance. Here, we use the term ‘transcript model’ to describe a distinct set of splice junctions paired with variable 5’ and 3’ ends. Bioinformatics software packages such as Kallisto use expectation-maximization to pseudo-align short reads to a transcriptome reference, generating abundance estimates for transcript and gene models^16^. These algorithms are effective in broadly identifying which transcripts the reads are compatible with, but they cannot tell exactly which ones were present. Long-distance contiguity is especially challenging. An additional drawback is that these methods depend heavily on the choice of the reference transcript annotation and, as such, they cannot identify novel transcript models. Another widely used approach to quantifying alternative splicing is to compute short read coverage of specific splice junctions or exons, and compare the resulting counts across samples using statistical tests^17,18^. While these methods are useful for detecting alternative exon usage, they do not overcome the fundamental limitations of short-read data with respect to assembling and assigning exactly which exons made up the source transcript.

Since 2012, third-generation sequencing platforms such as PacBio and Oxford Nanopore (ONT) have pioneered the use of long reads in genomics^19,20^. With read lengths of up to 60 kb for PacBio and up to 1 Mb for Oxford Nanopore, these reads can capture entire transcripts from end to end. They also offer the advantage of representing single molecules rather than amplified clusters, making them ideal for sequencing isoforms. Historically, the major drawbacks of long read technologies have been their relatively low throughput as well as high indel and mismatch error rates ranging up to 15-20%^19^. In the case of PacBio, these stochastic errors are mitigated by using circular consensus sequencing, in which multiple sequencing passes over the same molecule are used for error correction^21^. The exact error rate depands largely on the number of passes that a molecule receives. Computational methods have also been developed to correct errors in long reads, including hybrid approaches that incorporate short reads, and other methods that make use of reference annotations^22–25^.

Due to the low throughput of the original platforms, the conventional long-read transcriptome sequencing approach was to first catalog expressed isoforms using long reads from size-selected subsamples, and then map short reads to the resulting transcriptome references for the purpose of quantification^26–28^. PacBio popularized this method in mammals, plants, and beyond under the name “Iso-seq”. Recently, PacBio yields increased substantially, producing up to 8 million reads per SMRT cell on the Sequel 2 compared to 150,000 on the older RSII machines. Similar yield increases have been reported for Oxford Nanopore. This increased throughput has made direct long-read quantification more plausible. Unfortunately, most existing tools for analyzing long-read transcriptome data were not explicitly designed for this purpose. PacBio-affiliated software packages such as ICE-Quiver/Arrow and Cupcake ToFU generate *de novo* transcript models by clustering long reads and then merging them to generate one transcript model per cluster^26,29^. This is a particularly useful approach in species that lack a reference genome, but it comes with disadvantages. ICE-Quiver has been known to merge together transcripts from highly similar genes and can smooth over real differences of interest such as sequence variants and RNA editing events^30^. In addition, the algorithm is stochastic by nature, and cluster assignments for individual reads can vary substantially across different runs. Most existing programs for transcriptome-wide PacBio annotation and quantification rely on the ICE-Quiver or Cupcake ToFU outputs. For instance, SQANTI uses post-ToFU transcript models and their estimated abundances as the input to its annotation, quantification, and quality control pipeline^23^. Another set of pipelines such as FLAIR have been developed for analyzing Oxford Nanopore cDNA and direct RNA sequencing data^31^. As in ICE-Quiver, a common feature of these pipelines is the alignment of reads to each other before determining which known and novel transcripts are present.

Here, we present TALON, the ENCODE4 pipeline for simultaneous transcript discovery and quantification of long-read RNA-seq data regardless of platform. This pipeline is designed to explicitly track both known and novel transcripts across different bio-samples to allow for annotation and use of new isoforms. The full TALON pipeline is available on GitHub through the ENCODE4 Data Coordinating Center (DCC) at ENCODE-DCC/long-read-rna-pipeline and at mortazavilab/TALON. We first analyze the transcriptome of the GM12878 cell line using the PacBio and ONT to quantify the relative performance of both platforms. The TALON pipeline allows us to process PacBio and ONT data in a uniform fashion and make direct comparisons between the two. We evaluate the resulting transcriptomes relative to available CAGE, poly(A), and RNA-PET annotations in these cells and find that each long-read technology is affected by different artifacts. We then sequence the transcriptomes of adult mouse hippocampus and cortex to show the applicability of the TALON pipeline for the analysis of complex tissues. Overall, we demonstrate that current long-read platforms are suitable for quantifying and characterizing isoform-level expression of genes.

## RESULTS

### Tracking transcript novelty and quantification using TALON

To compare long read platforms side by side and to track isoforms consistently across multiple datasets, we developed a technology-agnostic long-read pipeline called TALON **(Figure 1a)**. This pipeline is designed to annotate full-length reads as known or novel transcripts and also to report the abundance for genes and transcripts across datasets. Starting from long reads mapped to the reference genome with a long-read aligner such as Minimap2, reference-based error correction is performed using TranscriptClean to remove microindels, mismatches, and noncanonical splice junctions in a variant-aware manner as previously described^25^. Noncanonical splice junctions are permitted in the final output only if they are supported by the splice annotation. Note that TALON expects reads to be oriented to the appropriate strand, which is typically achieved using platform-specific preprocessing in the case of cDNA reads **(Figure S1a-b).** After TranscriptClean, corrected reads are passed into the *talon_label_reads* TALON module, which records QC information for use by subsequent steps. In particular, long-read libraries built using poly(A) selection are prone to internal priming artifacts in A-rich regions of transcripts that result in truncated isoforms. Therefore, tracking the fraction of As following alignments is informative for TALON’s transcript filtering process. After the internal priming labels have been assigned, the reads are passed into the main *talon* module for annotation. In a talon run, each input SAM read is compared to known and previously observed novel transcript models on the basis of its splice junctions, start, and end points. This allows us to not only assign a novel gene or transcript identity where appropriate, but to track new transcript models and characterize how they differ from known ones. The result is a collection of all transcripts observed in each input dataset that can then be filtered, quantified, and compared using downstream TALON modules.

**Figure 1.**
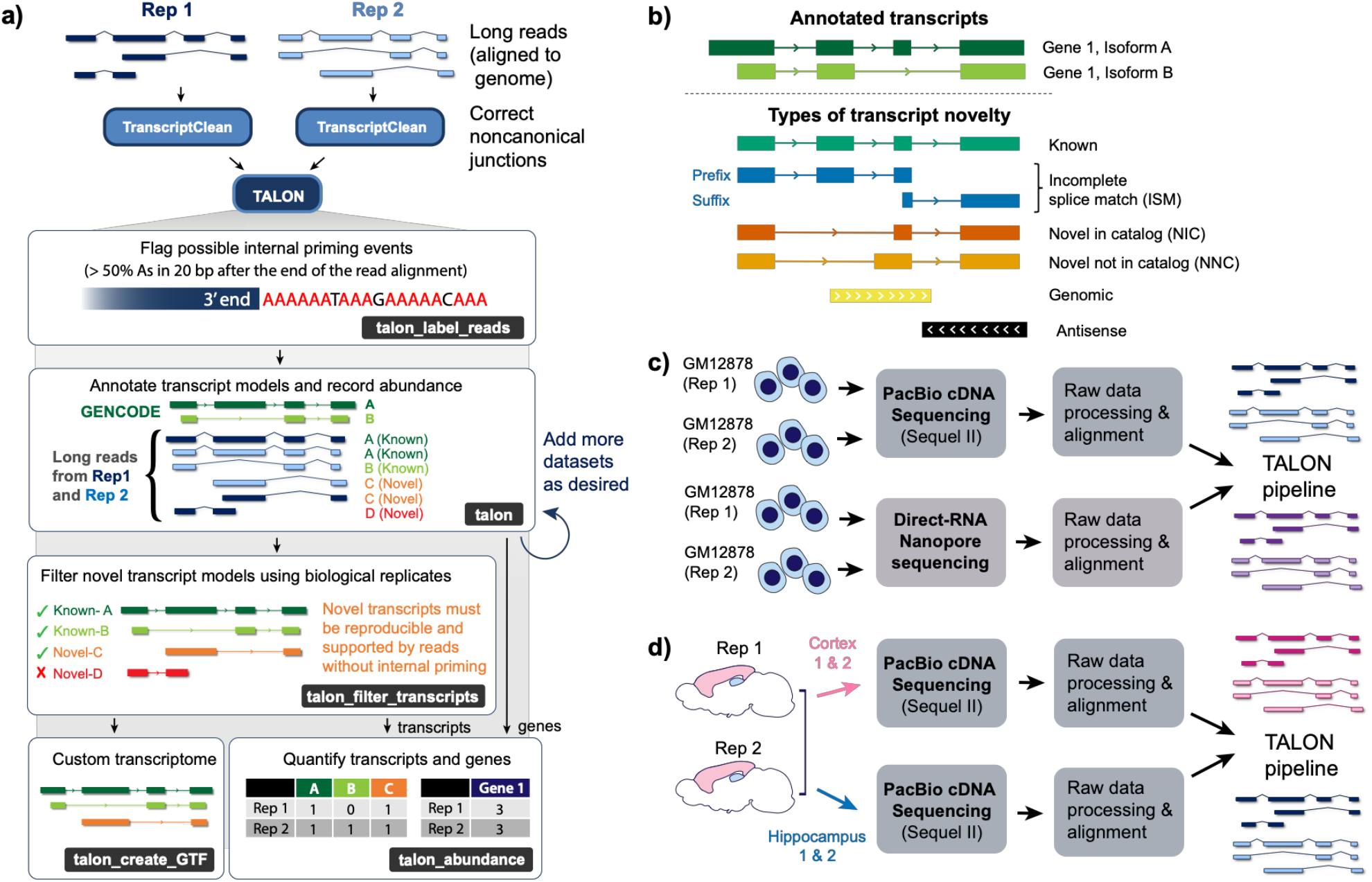
Overview of TALON. **a)** Long-read alignments from either technology are corrected with TranscriptClean for each biological replicate. Next, potential internal priming events are flagged by the talon_label_reads module. Labeled reads are passed into talon, where they are assigned a gene and transcript identity. The talon_abundance module computes gene expression directly from the talon results, whereas novel transcript models are filtered prior to quantification. Novel transcripts must be reproducibly detected n times in k datasets to pass the filter (default n = 5 and k = 2), and must not come from internally primed reads. **b)** Types of transcript novelty tracked by TALON. **c)** TALON can be used to compare different long-read sequencing technologies run on the same biological sample such as the human GM12878 cell line. **d)** TALON can also be used to compare genes and transcripts across different samples such as mouse hippocampus and cortex.

We adopted the nomenclature introduced by SQANTI to characterize the different types of transcript novelty in our datasets^23^. Query transcripts with splice junctions that perfectly match an existing model are deemed ‘known’ **(Figure 1b)**. Flexibility is allowed at the 5’ and 3’ ends. In cases where a transcript matches a subsection of a known transcript model and has a novel putative start or endpoint, it is considered an ‘incomplete splice match’ (ISM). TALON further subdivides the ISM category into prefix ISMs and suffix ISMs. The former refers to ISMs that match along the 5’ end of an existing transcript model, and the latter describes ISMs that match to the 3’ end. It is possible for a transcript to belong to more than one ISM category if it matches to different parts of several existing transcript models. The ISM category is useful as a means of quality control as libraries with a higher proportion of ISMs relative to known transcripts tend to be less than complete in terms of length and may harbor more artifacts. For instance, RNA degradation and incomplete reverse-transcription can lead to suffix ISMs. In Oxford Nanopore, pore blockages can produce suffix ISMs by prematurely stopping sequencing of the RNA. In the case of prefix ISMs, internal priming is the most likely culprit. However, not all ISMs are sequencing artifacts. To differentiate between a truly novel ISM transcript and one that is artifactual, it is useful to test against relevant orthogonal data such as CAGE, RNA-PET, or poly(A) annotations, which are often available from external databases. This can provide independent validation to support or reject a new 5’ or 3’ end seen in an ISM transcript.

The next category, novel in catalog (NIC), describes transcripts that have known splice donors and acceptors, but reveal new connections between them. This can be thought of as a novel arrangement of known exons. Novel not in catalog (NNC) transcripts contain at least one novel splice donor or acceptor, meaning that there is at least one novel exon boundary present. Genomic transcripts are either partial transcripts that do not share any splice junctions with overlapping genes or may come from DNA contamination in the samples, and are therefore discarded by the filter, reproducible or not. The antisense category consists of transcripts that overlap an existing gene, but are oriented in the opposite direction. If a transcript lacks any overlap with a known gene, then it is deemed intergenic. Taken together, the novelty categories allow us to examine the types of transcripts that we detect in our long-read datasets, to perform quality control, and to stratify or filter by category.

Biological replicates serve as an important means of verifying novel transcript discoveries. Although the accuracies of long-read platforms are improving, artifactual transcripts are still a problem, and may arise from a variety of technical sources. TALON streamlines the filtering process for multiple datasets by tracking transcript annotations and abundance in one place, where the information can be easily accessed and compared. Our filtering process uses the novelty labels assigned to each observed transcript model in order to remove likely artifacts. Observed transcripts that fully match counterparts in the GENCODE annotation are accepted immediately, but we require that novel transcripts be supported by at least 5 reads each in at least two biological replicate samples in order to be included in the downstream analysis. Furthermore, all five reads must all pass the internal priming cutoff (fraction As ≤ 0.5). These cutoffs can be adjusted by the user to accommodate different oligo-dT lengths or sequencing depths. As additional samples are sequenced, it is also possible to cross-reference novel transcripts across these datasets.

TALON quantification relies on the premise that each long read represents an individual transcript molecule sequenced. This allows us to quantify expression by simply counting the number of individual reads that were assigned to a particular transcript or gene and then converting these values into units of transcripts per million (TPM) to adjust for library size. For gene-level expression values, we include all reads assigned to a locus in the computation, since even incomplete transcripts (ISMs) that did not meet the threshold to become a new transcript model are informative for the overall gene expression level. On the transcript level, however, we apply the TALON filters in order to avoid quantifying transcript models with insufficient evidence.

To demonstrate the utility of TALON, we applied it in two different settings **(Table S1)**. First, we compared long-read GM12878 data sequenced on different platforms: PacBio Sequel II and direct-RNA ONT **(Figure 1c; Table S3).** Then, we used TALON to analyze gene and isoform-level expression across the complex tissues of cortex and hippocampus in mouse **(Figure 1d; Table S3)**. In each case, we sequenced at least 6 million raw reads per replicate. Spike-in RNA variants (SIRVs) in our samples provided us with an opportunity to evaluate TALON filtering on artificial sequences with fully known splice patterns **(Table S4-5)**. The expected outcome in an error-free setting would be to detect exactly 69 known isoforms from a total of 7 SIRV genes, and to detect zero novelty. After applying the TALON transcript filter (including the internal priming cutoff) to SIRVs sequenced with the two PacBio GM12878 replicates, we detected 67 known SIRV transcripts and only 13 novel models **(Figure S2a)**. 96% of the filtered reads matched a known isoform **(Figure S2b)**, In contrast, the unfiltered SIRV data contained a much higher fraction of artifactual novel transcripts **(Figure S2c,d)**. The ISM category was the most common form of novelty, accounting for between 5 and 6% of the unfiltered reads by replicate. About 60% of the reads assigned to prefix ISMs displayed evidence of internal priming, suggesting that this is a substantial artifact of cDNA sequencing in PacBio **(Figure S2e)**. The TALON filter was highly effective in removing these transcripts-after filtering, only 9 ISM models remained. Overall, these results indicate that the TALON filter is effective at removing artifactual transcript models.

### Performance of TALON on human ENCODE Tier 1 PacBio data

We then turned our attention to applying TALON to GM12878 reads mapped onto the human genome **(Table S6-S8)**. TALON detected 15,727 known GENCODE genes and 26,841 GENCODE transcripts in GM12878 across the two replicates. The number of known genes is smaller than the number of known transcripts because known genes can be detected through novel transcripts as well as known ones. The analysis also called 359 unknown gene models, the majority of which consisted of monoexonic transcripts mapped as antisense within a known gene locus. The TALON N50 read lengths for Rep 1 and Rep 2 were 1,877 and 1,791 nucleotides, respectively, which is in line with the expected length distribution of most mammalian mRNA transcripts **(Fig S3)**.

We next computed the expression level for each known GENCODE gene across the PacBio data. For this quantification, we included all long reads assigned to a locus in these counts because even incomplete transcripts are informative for the overall gene expression level. The resulting gene expression levels were highly correlated across biological PacBio replicates of each cell line (Pearson r = 0.97, Spearman rho = 0.92) **(Figure 2a)**. This shows that our PacBio primary data coupled with the TALON pipeline produces reproducible quantifications of gene expression.

**Figure 2.**
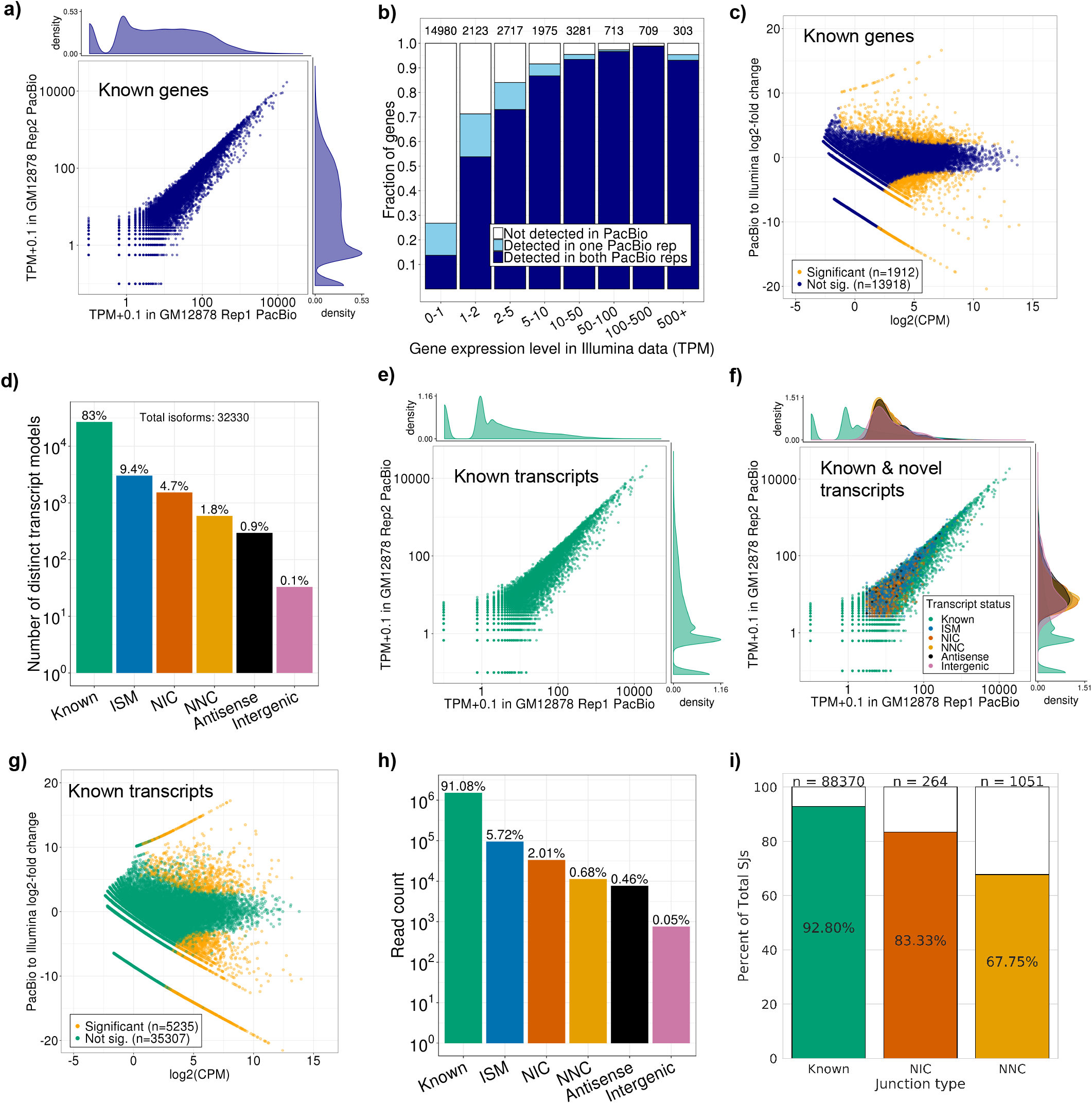
Performance of TALON on PacBio transcripts from GM12878 cell line. **a)** Expression level of known genes (GENCODE v29) in each biological replicate (Pearson r = 0.97, Spearman rho = 0.92). **b)** Proportion of genes expressed in Illumina RNA-seq data of GM12878 that are also detected in PacBio, binned by Illumina expression level (TPM). **c)** Comparison of gene expression levels for known genes in the PacBio and Illumina RNA-seq platforms. **d)** Number of distinct transcript isoforms observed in each novelty category. **e)** Expression level of known transcript models in each biological replicate (Pearson r = 0.97, Spearman rho = 0.73). **f)** Expression of transcript models in each biological replicate, labeled by their novelty assignments (Pearson r = 0.97, Spearman rho = 0.83). **g)** Comparison of known transcript expression levels in the PacBio and Illumina RNA-seq platforms. **h)** Total number of PacBio reads assigned to each novelty category after transcript filtering (Rep 1). **i)** Percentage of known and novel PacBio GM12878 splice junctions supported by Illumina. Junctions labeled NIC indicate novel combinations of known splice sites, while NNC junctions included a new donor and/or acceptor.

We also compared our PacBio results to short-read RNA-seq data from the same cell line. First, we examined how often PacBio was able to detect known genes as a function of their short-read expression level **(Figure 2b)**. As expected, genes at the lower range of expression (< 2 TPM from short reads) were less likely to be detected by PacBio, but upwards of 70% of genes expressed above 2 TPM were reproducibly detected. Overall, the expression levels of the 14,947 genes detectable in both PacBio and Illumina correlated well across platforms (Spearman rho 0.78). We conducted a differential expression analysis to further examine how much gene expression levels vary between the platforms. The log fold change between PacBio and Illumina was computed using the exact test method in EdgeR, and Bonferroni correction for multiple testing was performed on the resulting p-values **(Table S9)**. This analysis revealed that there was no significant difference in expression levels for most genes **(Figure 2c).** However, a subset of genes showed significant fold change differences, including 773 that were higher in PacBio and 1,139 that were higher in Illumina. Genes expressed significantly higher in Illumina tended to have longer median transcript lengths on average than those that were not differentially expressed or that were expressed more highly in PacBio **(Figure S4a)**. This suggests that these PacBio data under-detect the longest transcripts (greater than 5 kb) when no size selection is applied. Genes with higher expression in PacBio had significantly higher median GC content as a group (adjusted p = 4.950e-08) than those that were higher in Illumina (**Figure S4b)**. It is possible that this is related to the GC bias known to affect Illumina next-generation sequencing. Overall, non-size selected PacBio libraries detect most of the genes expressed at 1 or more TPM in Illumina.

Having established that TALON can quantify gene-level expression on the basis of long reads, we moved on to transcript-level quantification. As expected, most of the transcript models identified in our analysis of the extensively-studied GM12878 cell line were known matches to the GENCODE annotation **(Figure 2d)**. The expression levels of detected known transcripts were highly correlated across PacBio biological replicates (Pearson r = 0.97, Spearman rho = 0.73) **(Figure 2e)**. Novel transcript models displayed even stronger expression correlations, likely related to the stringent abundance and filtering requirements that were applied to them (Pearson r = 0.97, Spearman rho = 0.83) **(Figure 2f)**. PacBio transcript expression levels were not significantly different for 87% of GENCODE transcripts when compared to short-read expression levels **(Figure 2g; Table S10)**. The known, NIC, and NNC isoform categories account for about 94% of the filtered PacBio reads, with known transcripts making up 91.1% of the reads **(Figure 2h)**. NIC and NNC transcripts contained a larger number of exons on average than the other novelty categories, and also tended to come from longer reads **(Figure S5a-b)**. To evaluate the canonical junctions found in the PacBio reads, we compared them to junctions called from the short-read Illumina GM12878 RNA-seq data using STAR^32^. 83% of novel PacBio splice junctions featuring known splice donors/acceptors had short-read support **(Figure 2i)**. The majority of PacBio junctions with a novel splice donor and/or acceptor were supported as well. Overall, these results indicate that we can reliably annotate and quantify transcript models using our long-read pipeline.

GM12878 is an Epstein-Barr Virus (EBV) transformed lymphoblastoid cell line (LCL). We were therefore able to analyze the gene and transcript expression of EBV within the GM12878 PacBio transcriptome. We found that EBV transcripts are detectable using long-read sequencing, and that these transcripts can be quantified, annotated, and assessed for their novelty using TALON **(Table S11-13)**. Overall, 25 known and 4 post-filter novel EBV transcript isoforms were detected and 28 known EBV genes were detected **(Figure S6a-b)**. Many detected transcripts belong to the *EBNA* gene family **(Figure S6c)**, which code for proteins that are essential to the virus’ ability to transform infected cells into LCLs ^33^, and are typically among the most highly expressed genes from the EBV chromosome in LCLs.^34^ Consistent with the novel transcript models detected by TALON, the *EBNA* transcripts have previously been identified as heavily alternatively spliced^35^.

### Performance of TALON on Oxford Nanopore data and comparison with PacBio

Oxford Nanopore is an alternative long-read sequencing platform that offers the option of direct RNA sequencing ^36^. While the protocol involves one reverse-transcription step, this is primarily for the purpose of removing secondary RNA structure and ultimately only the RNA strand is sequenced. In order to demonstrate the applicability of TALON to the Nanopore platform, we directly sequenced RNA from two biological replicates of GM12878 to a depth of at least 2 million basecalled reads per replicate. After alignment with Minimap2^37^, each replicate was processed with the TALON pipeline as described for PacBio **(Table S14-16)**. The TALON N50 read lengths for the datasets were 1,269 nucleotides for Rep 1 and 989 for Rep 2 **(Fig S7a-b)**. Although the starting number of reads was lower than in our PacBio transcriptomes, we detected ~13,500 known GENCODE genes and ~18,000 known isoforms in GM12878. Gene and transcript expression levels across the two GM12878 ONT replicates correlated with each other (gene Pearson r = 0.99, gene Spearman rho = 0.92; known transcript Pearson r = 0.97, known transcript Spearman rho = 0.64) **(Figure 3a-b)**. When we labeled the transcripts by their novelty type, it became apparent that differences in isoform-level expression between ONT replicates are largely driven by overrepresentation of novel ISM transcript models **(Figure 3c-d)**. This leads us to believe that ONT is more sensitive to degradation events or is prone to stopping mid-transcript during sequencing, which may explain the high ISM numbers in our data.

**Figure 3.**
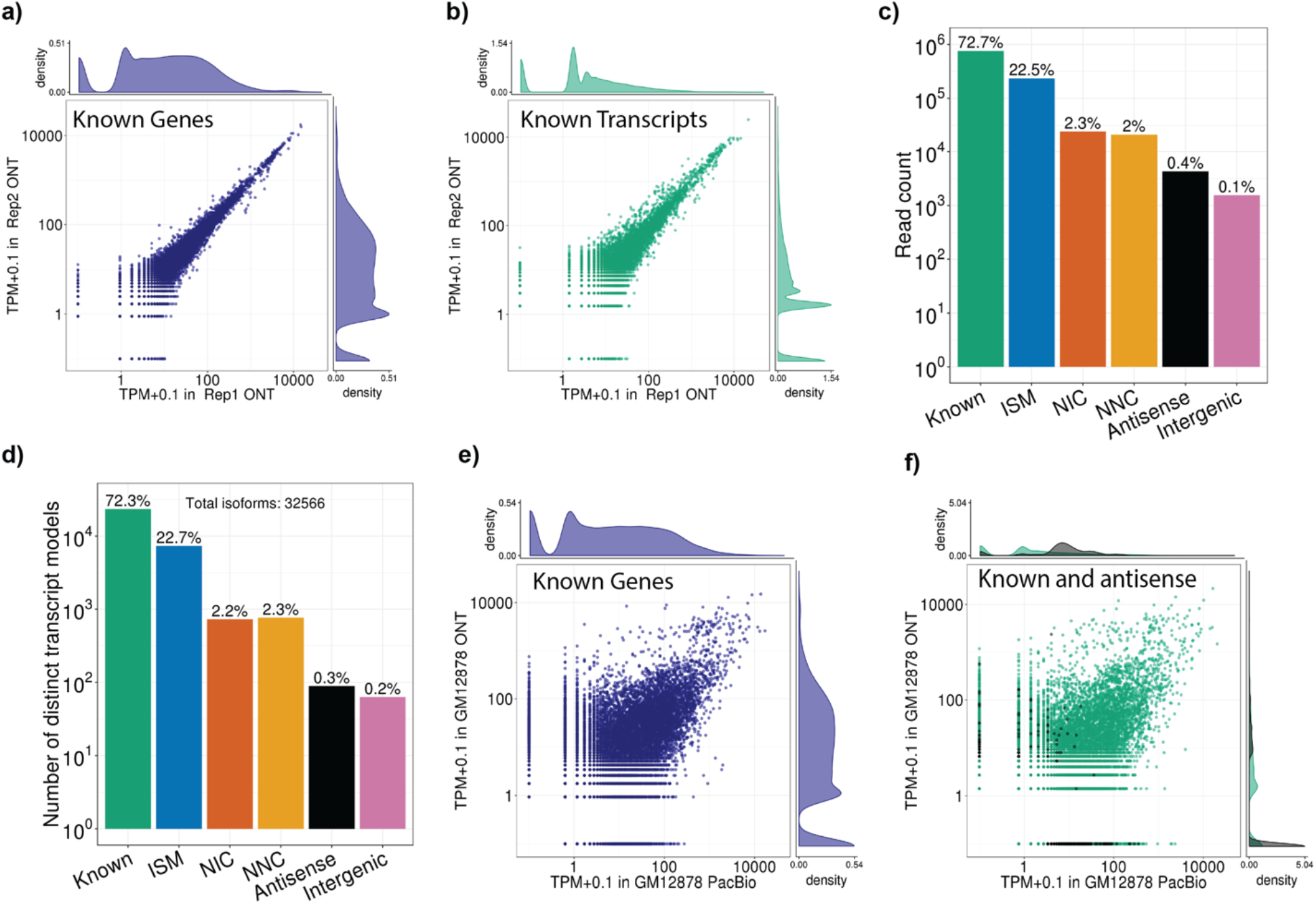
Comparison of Oxford Nanopore direct RNA-seq transcriptome with Pacbio transcriptome in GM12878. **a-b)** 2 GM12878 replicates were sequenced using the MinIon platform and analyzed using TALON pipeline to measure **a)** gene expression (Pearson r = 0.99, Spearman rho = 0.92) and **b)** transcript expression (Pearson r = 0.97, Spearman rho = 0.64). **c)** Total read count per novelty category. There is a substantially larger fraction of ISM reads than full-length known compared to PacBio (Fig 2h). **d)** Number of distinct isoforms by novelty category. **e-f)** Correlations between ONT direct RNA-seq and PacBio with respect to **e)** gene expression (Pearson r = 0.58, Spearman rho = 0.63) and **f)** transcript expression (Pearson r = 0.5, Spearman rho = 0.18).

Next, we compared gene and transcript expression levels across the PacBio and ONT platforms in GM12878 **(Figure 3e; Table S17-19)**. These were well-correlated at the gene level, but there were interesting differences at the transcript level. For instance, ISMs were overrepresented in ONT relative to PacBio, suggesting that the former had more difficulty sequencing full-length transcripts (**Fig S7c**). On the other hand, of 414 total antisense transcripts called across the platforms, 276 were unique to PacBio, whereas only 26 were detected in ONT alone **(Figure 3f)**. This likely means that the majority of antisense transcripts were in fact artifacts of the reverse transcription steps required for PacBio, demonstrating a drawback of conversion to cDNA before sequencing, at least by the standard methods used for PacBio. Interestingly, there is a set of 88 genes with TPM > 10 in both technologies that are detected as more than 10-fold more highly expressed in Oxford Nanopore **(Table S20-21)**, which could represent further under-representation of these transcripts due to reverse transcription biases. Among the genes enriched in Oxford Nanopore we found a subset related to mitochondrial functions (MT-RNR1, MTCO1, MT-CO2, MT-ATP6, MT-CO3 and MT-CYB), that have been previously characterized as a benchmark for direct RNA-seq performance as pointed out by other groups^31^. Although some mitochondrial genes are subject to a deadenylation process, mature mt-mRNA transcripts contain a non-templated sequence of poly(A)s^38^. This fact, along with the minimal processing steps before sequencing, might explain the higher detection levels of these genes on the Oxford Nanopore platform compared to PacBio.

### Comparison of TALON and FLAIR on GM12878 Pacbio and ONT data

FLAIR is another recent pipeline designed to identify and quantify transcripts in long-read PacBio or ONT data^39^. To compare FLAIR and TALON, we ran FLAIR on the full-length, non-chimeric PacBio reads from GM12878 replicates 1 and 2 as described in the Supplementary Methods. We then compared the FLAIR quantification results to those generated by TALON in Figure 2. Similarly to TALON, FLAIR reported strong gene and transcript-level expression correlations across biological replicates (FLAIR Pearson r = 0.96, Spearman rho = 0.94 for known genes and Pearson r = 0.96 Spearman rho = 0.88 for known transcripts in GM12878). However, FLAIR was less sensitive than TALON with respect to detecting known genes and transcripts **(Table S22-23)**. For instance, in GM12878, TALON detected 2,525 more GENCODE genes than FLAIR that were also expressed in the corresponding short-read data **(Figure S8a)**. Recognizing that FLAIR was initially developed for ONT data, we ran the same comparison on our direct-RNA ONT GM12878 datasets **(Fig. S8b)**. As in the PacBio analysis, FLAIR detected fewer known genes and transcripts in the ONT data than TALON **(Table S22-23)**. This discrepancy was particularly pronounced at lower expression levels, but applied to genes with > 50 TPM in Illumina as well (**Fig. S8b).** Taken together, these results demonstrate that TALON is currently more sensitive to known genes and transcripts than FLAIR in the same datasets.

### Assessing completeness of TALON transcript models using CAGE, poly(A) motifs, and RNA-PET

The exonuclease treatment of our samples at the RNA stage and the full-length classification step *in silico* are intended to ensure that the transcripts at the end of our pipeline have intact 5’ and 3’ ends. To verify completeness, we performed an integrative analysis comparing our TALON transcript models with data from the CAGE and RNA-PET assays, as well as computationally identified poly(A) motifs. For known transcript models, the annotated GENCODE 5’ and 3’ sites were used.

CAGE is a genome-wide method of annotating transcription start sites that works by trapping the 5’ end cap of a mature mRNA transcript using an antibody and then sequencing its 5’ end. To validate the 5’ ends of our long-read transcript models, we compared them to CAGE-derived TSSs from the FANTOM5 project. 76% of known GENCODE transcripts in our GM12878 PacBio transcriptome had CAGE support **(Figure 4a)**. Transcripts in the prefix ISM category were overwhelmingly supported (97%), whereas suffix ISMs were not (34%). 94% of NIC and 87% of NNC transcripts were supported by CAGE, indicating that their 5’ ends were at least as reliable as those of the known transcripts. However, the antisense PacBio transcripts had scant support, lending credence to the idea that they are largely reverse-transcription artifacts. We observed similar CAGE trends in our ONT transcriptome **(Figure 4b)**, although notably, most transcript categories tended to have lower rates of support than in the corresponding PacBio transcriptome.

**Figure 4.**
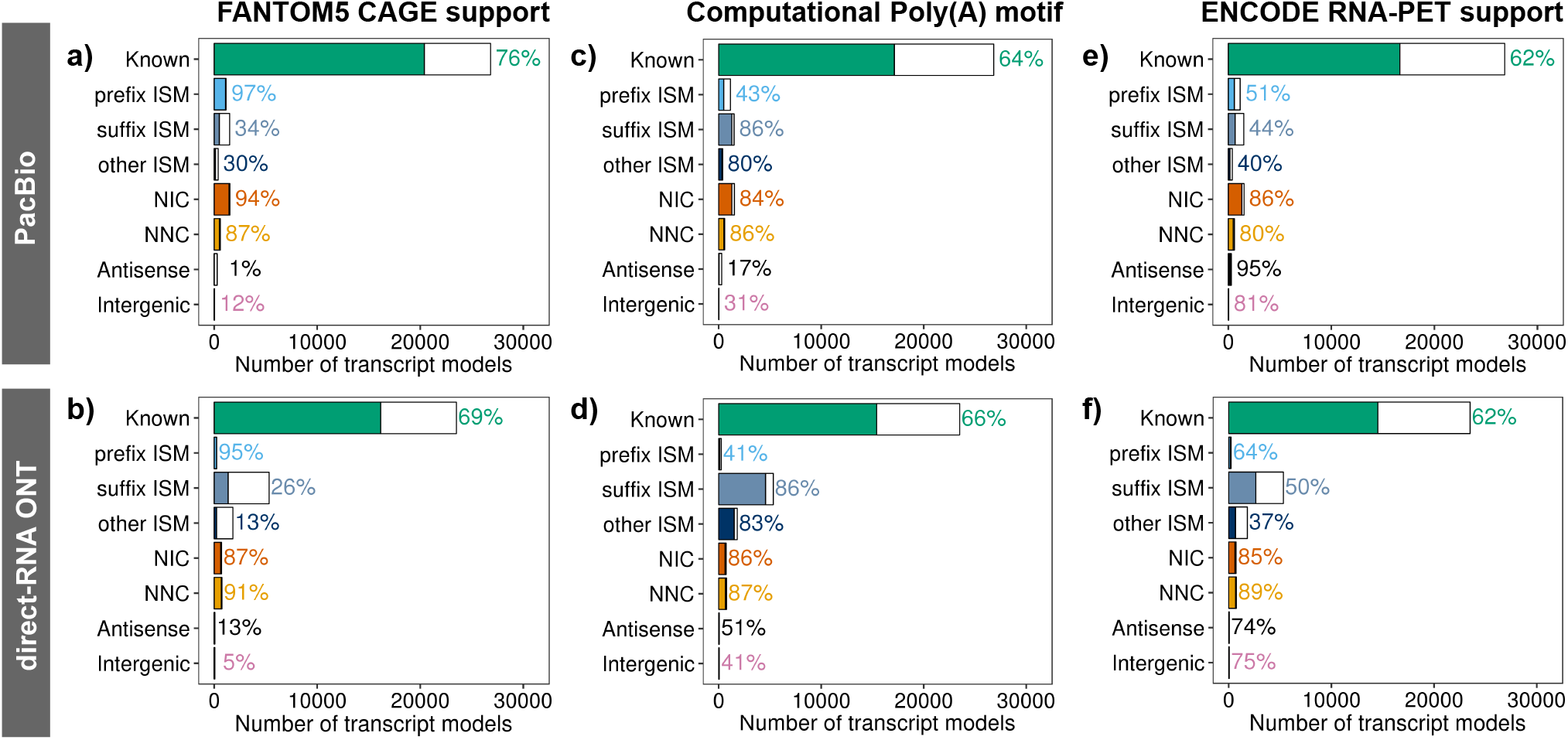
External validation of transcript model ends by novelty category. **a)** Percentage of TALON transcript models with CAGE support for their 5’ end by novelty category (GM12878 PacBio). **b** Percentage of TALON transcript models with a poly(A) motif identified at their 3’ end (GM12878 PacBio). **c)** Percentage of TALON transcript models with RNA-PET support for their 5’-3’ end pair (GM12878 PacBio). **d)** Percentage of TALON transcript models with CAGE support for their 5’ end by novelty category (GM12878 ONT). **e)** Percentage of TALON transcript models with a poly(A) motif identified at their 3’ end (GM12878 ONT). **f)** Percentage of TALON transcript models with RNA-PET support for their 5’-3’ end pair (GM12878 ONT).

To examine transcript completeness at the 3’ end, we conducted a computational poly(A) motif analysis of our long-read transcript models. This entailed scanning the last 35 bases of each transcript sequence to look for the presence of a known poly(A) motif. In PacBio, 64% of known transcripts contained such a motif **(Figure 4c)**. Rates of support were also high in the suffix ISM, other ISM, NIC, and NNC categories (86%, 80%, 84%, and 86% respectively). As expected, only 43% of the prefix ISMs contained a poly(A) motif, indicating that many of these transcripts may be artifactual. Overall, similar trends were observed in the ONT transcripts **(Figure 4d)**.

Finally, we sought to validate the 5’-3’ pairings in our transcript models using publicly available RNA-PET data from the ENCODE consortium for both PacBio and ONT transcriptomes **(Figure 4e-f)**. This assay marks the start and endpoints of individual cDNA transcripts by circularizing and sequencing them with paired-end tags. This data type was lower-throughput than the more recently generated CAGE data, which helps explain the lower rates of RNA-PET support for known transcripts. We nevertheless observed strong RNA-PET support for NIC and NNC transcripts in both PacBio and Oxford Nanopore. Of the three ISM categories, prefix ISMs were the most likely to have RNA-PET support for their 5’-3’ end pairing. Antisense transcripts had extremely high rates of RNA-PET support. The RNA-PET protocol uses reverse transcription, and therefore it is possible that this assay is prone to the same types of antisense artifacts as PacBio.

Taken together, the results of our CAGE, poly(A), and RNA-PET analyses indicated that most NIC and NNC transcript models derived from long reads have intact 5’ and 3’ ends, which argues that they represent full-length RNA’s. However, inferred transcripts in the ISM novelty category require more scrutiny. As expected based on the category definition, prefix ISMs had reliable 5’ sites, but their 3’ ends were potentially incomplete in many cases. The reverse was true for suffix ISMs. In both cases, this suggests that many are technical artifacts. In general, the PacBio platform did a better job of capturing complete transcripts in our hands than did direct-RNA ONT, and offered the additional benefit of higher throughput.

### Comparison of PacBio transcriptomes of mouse cortex and hippocampus

After testing and characterizing TALON on PacBio data in a homogeneous cell line, we applied it to begin to discover and quantify isoforms in the complex brain regions of the mouse cortex and hippocampus. The cortex and hippocampus are critical regions of the brain for learning because of their functions of neural integration and memory, respectively^41^. Therefore, these regions have been characterized extensively under different conditions and models in order to understand their gene expression profiles^42^. These brain regions are much more complex in cell type composition than isolated cell lines, and the two regions have both similar and disctint cell types. Regulation of cell type diversity is key during their development, aging, and in disease, with both known and likely undiscovered differences in gene and isoform-level expression^42^. In addition, the brain at large is known to have a high alternative splicing ratio when compared to other tissues ^40^.

We sequenced two PacBio Sequel II replicates each of cortex and hippocampus to a minimum depth of 6 million raw reads per replicate and ran TALON on them **(Table S24-26)**. Gene expression was highly correlated across biological replicates (cortex Pearson r = 0.96, Spearman rho = 0.95 and hippocampus Pearson r = 0.89 Spearman rho = 0.83) as was transcript-level expression (cortex Spearman rho = 0.84 and hippocampus Spearman rho = 0.73) **(Fig. S9a-d)**. On average, we detected 17,000 known genes and 26,000 known transcripts for each tissue. The diversity of the isoform novelty categories was similar between cortex and hippocampus **(Figure 5a-b)**. We identified 694 differentially expressed transcripts isoforms from a total of 612 genes (log fold-change > 1 and adjusted pvalue < 0.01), including 607 known and 87 novel transcript models **(Figure 5c; Table S27)**. This included differences between known transcripts for genes such as Pnisr, which is a splicing factor involved in aging^43^. Other examples involving novel isoforms include the maternally expressed 3 (Meg3) long-coding RNA gene for which we detected two NIC transcript models that were enriched in cortex. Meg3 is thought to be involved in controlling vascularization in the brain by inhibiting angiogenesis^44^ and is highly expressed. In addition, an NNC transcript of Amigo2 was selectively enriched in hippocampus. Amigo2 is known to be upregulated in the CA2 and CA3a regions^45^ and is also commonly used as a marker of astrocyte activation^46^.

**Figure 5.**
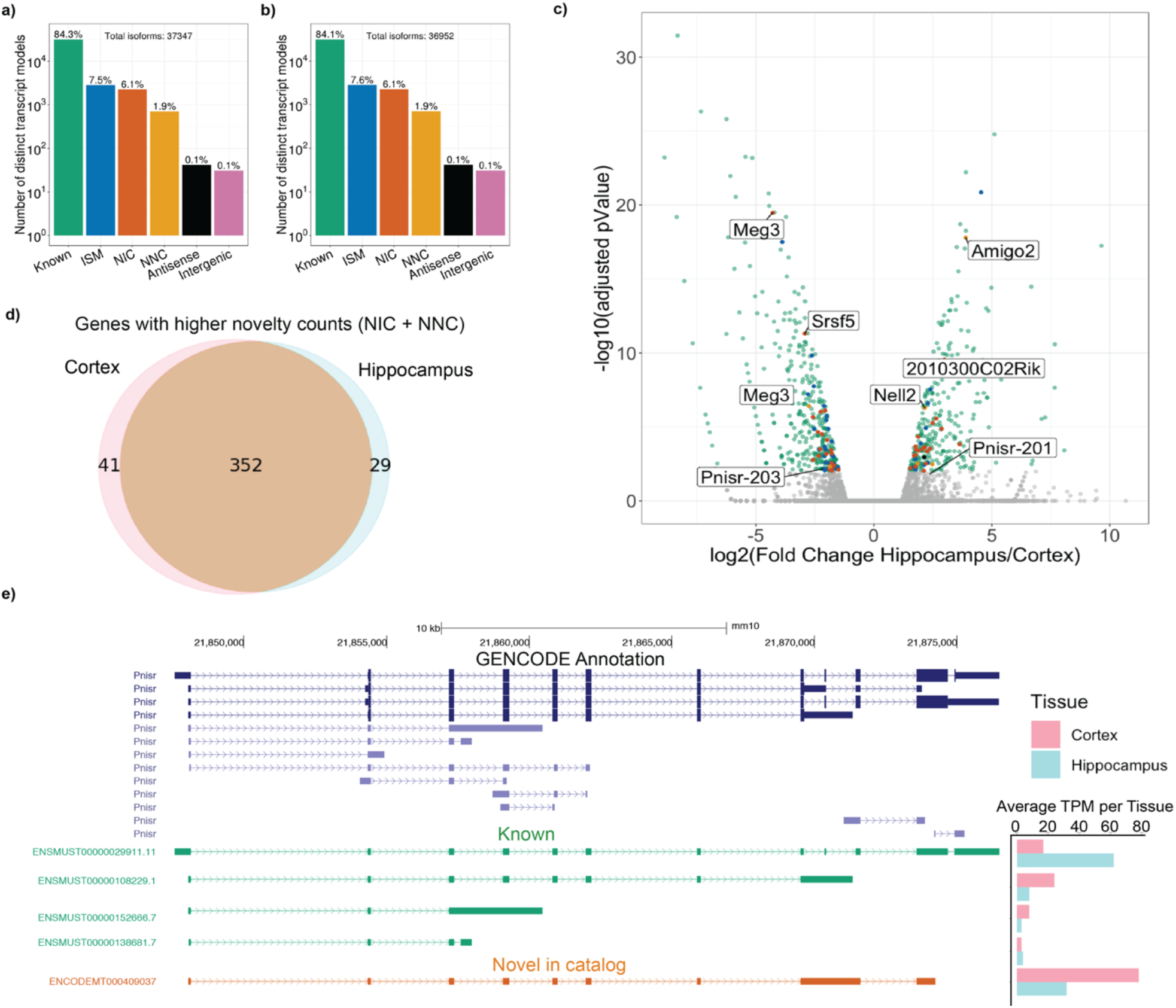
PacBio transcriptomes of 8-month male adult mouse cortex and hippocampus. Novelty assignments of distinct transcript models detected in one representative replicate each of **a)** cortex and **b)** hippocampus. **c)** Differential isoform expression in hippocampus and cortex. Transcripts with a fold change > 1 and an adjusted p value of < 0.01 are colored according to their novelty status and labeled with their corresponding gene name (color scheme in panels a,b) Detection of genes with greater novel read counts (NIC + NNC) than known. **e)** Pnisr UCSC genome browser visualization showing a new combination of exons detected by PacBio. Expression levels of each isoform detected are is plotted on the right for cortex and hippocampus.

We extended our transcript analysis by asking whether there are specific sets of genes with more novel transcript expression than known transcript expression. Specifically, we focused on genes that had more reads assigned to NIC and NNC novel transcript isoforms than known transcript isoforms in either brain region and found a shared set of 352 genes with an additional 29 and 41 that were specific to cortex and hippocampus respectively **(Figure 5d; Table S28).** This includes an NIC isoform of Pnisr (ENCODEMT000904037) that is enriched in the cortex **(Figure 5e)**. These analyses show that full-length transcriptome sequencing can detect isoform differences even in intensively studied tissues and cell types. These differences would be difficult to recover from short reads alone, especially for the NIC isoforms.

## DISCUSSION

We demonstrated here that with sufficient sequencing depth, long read data can be used to reproducibly quantify gene and transcript expression in homogeneous cell lines and in complex tissues. Our technology-agnostic long-read pipeline, TALON, simplifies the process of comparing long-read transcriptomes across different datasets and allowed PacBio and ONT transcriptomes to be directly compared. We found that our PacBio results are reasonably well-correlated with Illumina short-read data, particularly for gene expression levels above 2 TPM. We also found that, in our hands, the current PacBio platform captured more complete transcript models than did the current direct-RNA ONT, but that the former is prone to antisense transcript artifacts that apparently stem from the reverse transcription step into cDNA. It is also likely that many of the ISM-class of transcripts that we detected more prominently in ONT are false positives due to the pore ceasing to sequence midway through an RNA. While over 80% of the transcript models we detect in the well-studied GM12878 cell line were already known, we nonetheless found evidence of a number of new NIC and NNC transcript models that are supported independently by 5’ and 3’ ends from other genomics assays.

In contrast to the homogeneous and frequently-studied GM12878 cell line, we found that a substantial number of genes in the mouse cortex and hippocampus produced more novel (NIC and NNC) isoform reads than known isoforms. Not surprisingly, this suggests that we are still underestimating the overall contributions of alternative splicing in tissues that are both more complex in terms of cell composition and also less comprehensively measured to date. At this time, the goal of producing a reference-level annotation transcriptome for any given cell type or tissue is well-served by the PacBio platform, but our results also make it clear that any platform that provides transcript information by direct RNA sequencing, as the RNA ONT platform now does, makes a different and important contribution. At our current PacBio sequencing depth, we do not expect to encounter substantial issues with lack of complexity in our bulk cDNA libraries. However, as long-read cDNA sequencing depths increase, reads from PCR duplicates may become much more prevalent and would be difficult to detect without UMIs. This is never an issue with direct-RNA sequencing on the Nanopore platform because each read must correspond to a distinct mRNA molecule. As iterative advances are made on these platforms, and as other long-read systems are added, the ability to process and compare the outputs from all versions of all systems in a platform-agnostic way will be increasingly important.

In addition to the technology-specific challenges of each long-read platform, we identified some shared issues. While both PacBio and ONT could sequence most genes expressed in the cells, some very long transcripts were conspicuously missing or under-represented in our data. For instance, in GM12878, we only detected 3 reads that fully matched known isoforms of the highly expressed XIST gene in terms of their splice junctions. Even the longest of these reads (3,539 nt) was missing several kb from the 5’ and 3’ ends of the annotated GENCODE model. More generally, while NIC and NNC transcript models looked identical to or better than known transcripts in terms of CAGE, poly(A), and RNA-PET validation, ISMs represent a challenge for both technologies. This is particularly pressing as we detect more such ISMs in our brain tissue biosamples than in cell lines. We expect that parsing ISMs will be a challenge in human post-mortem tissue samples, including reference collection efforts for ENCODE4, because RNA quality is typically lower than what we obtained from cell lines and fresh mouse tissue sources. The “Iso-seq” approach, which intentionally enriches long-read size catagories, has been to collapse ISM reads onto known transcripts. However, our results show that a subset of ISMs do have independent CAGE and 3’ end support. Thus, biological ISM forms are difficult to distinguish from truncated reads without, at minimum, some independent CAGE support. Interestingly, the XIST locus is crowded with CAGE peaks throughout its longest transcript model, suggesting that there may be multiple “shorter” isoforms produced than previously appreciated, with evidence for them having been ignored due to the lack of resolution using short-reads alone. ISMs are, in any case, useful models to incorporate into gene expression quantifications. With additional datasets and evidence for training, we anticipate that machine learning techniques will allow us to discriminate real ISMs from technical artifacts. Until then, it seems prudent to ignore ISMs for transcript discovery in the absence of CAGE (or similar) support.

Clear challenges remain to generate fully comprehensive, high-fidelity long-read transcriptomes because of the still relatively noisy sequencing methods and imperfectly preserved RNAs. That said, our results show that current long-read methods are already demonstrably superior to “pooled” short-read RNA-seq for reference annotation-level transcriptomics if high quality mRNA can be extracted. The resulting gain in clarity with respect to long-range isoform structure and associated isoform-specific quantification is already substantial, although relatively high costs remain a limiting factor. At the time of this study, our long-read data costs were roughly an order of magnitude higher than their short-read counterpart, although a useful perspective is that this cost is comparable to that of short-read RNA-seq 10 years ago. We expect that long-read data will decrease similarly in cost per experiment as these platforms mature. Even in the domain of single-cell RNA-seq, which is currently thriving on short single-reads for molecule counting, long-read formats are beginning to be applied, aiming to capture the richness of isoform variation and regulation on a per-cell and per-cell-type basis^47^. That said, short-read transcriptomes will surely continue to play a prominent role for short RNA class substrates, for intractably degraded RNAs, and, increasingly, in biological settings where a few long-read transcriptomes can provide a reference against which larger numbers of companion short-read samples can be quantified. Ultimately, the transition to routine long-read transcriptome quantification will allow biologists to achieve clarity about functional mRNA isoform choices and their inferred protein products for any cell type, tissue, or disease state.

## METHODS

### Sample collection and RNA extraction

GM12878 cells were grown and harvested as described in the ENCODE consortium protocols (encodeproject.org). Total RNA was extracted using the QIAGEN RNAEasy Plus kit (Cat. No. 74134). All animal experimental procedures were approved by the Institutional Animal Care and Use Committee of University of California, Irvine, and performed in accordance with the NIH Guide for the Care and Use of Laboratory Animals.

Mice were anesthetized with CO_2_ and perfused with phosphate buffered saline (PBS) for 5-7 minutes until most organs are clean from blood. Hippocampus and Cortex from two 8-month male C57BL/6 mice were dissected and collected in HBSS no calcium no magnesium solution (cat. No. 14170112). Tissues were homogenized using the QIAshredder while in lysis buffer included in the QIAGEN RNAEasy Plus kit (Cat. No. 74134). Total RNA extraction was done following the vendor instructions. To degrade mRNA without a 5’ cap, total RNA was exposed to an exonuclease treatment using Terminator™ 5’-Phosphate-Dependent Exonuclease (Cat. No. TER51020).

### PacBio library preparation, sequencing, and initial data processing

Starting from the depleted RNA, we followed a modified version of the SMART-seq2 protocol to synthesize cDNA^48^. 1000 ng of cDNA were used as input for the PacBio library prep following the SMRTbell Template Prep Kit 2.0 instructions. Sequencing was done on the PacBio Sequel II machine, allocating 1 SMRT cell per biological replicate. Raw PacBio subreads were processed into circular consensus reads using the Circular Consensus step (CCS v4.0.0) from the SMRTanalysis 8.0 software suite (parameters: --noPolish --minLength=10 --minPasses=3 –min-rq=0.9 – min-snr=2.5) **(Figure S1a)**. Next, adapter configurations were identified and removed using Lima v1.10.0 (parameters: **--isoseq --num-threads 12 --min-score 0 --min-end-score 0 --min-signal-increase 10 --min-score-lead 0). After this,** full-length non-chimeric (FLNC) reads were extracted using the Isoseq3 Refine step (v3.2.2; parameters: **--min-polya-length 20 --require-polya)**. This program considers a read to be FLNC if it contained the expected arrangement of 5’ and 3’ PacBio primers at the Lima stage as well as a poly-(A) tail. Refine orients the reads to the correct strand (reverse-complementing sequences as necessary), and removes the poly-A tails. The resulting FLNC reads were mapped to the reference genome using Minimap2 version 2.17 (GRCh38 assembly for human cell types, and mm10 for mouse) with parameters recommended by the Minimap2 documentation for PacBio (-ax splice:hq -uf --MD).

### Illumina library preparation and sequencing for mouse brain samples

Starting from the same cDNA used for the mouse brain PacBio libraries, we built short-read libraries using the Nextera DNA Flex Library Prep Kit (https://www.illumina.com/products/by-type/sequencing-kits/library-prep-kits/nextera-dna-flex.html?scid=2017249vu1 <%22>). These libraries were sequenced on an Illumina NextSeq500 to a minimum of 50 million paired-75bp reads per sample.

### ONT library preparation and sequencing

Starting from 45 μg of depleted RNA, we proceeded to the direct-RNA library prep following the RNA-002 kit instructions. Reverse transcription was used to get rid of secondary RNA structures. We used R9.4 flowcells and MinKNOWN 2.0 was used to run the samples until having 2 millionraw reads. Basecalling was performed on the direct RNA ONT reads using ONT Guppy 3.2.1+334123b (parameters: -r --flowcell FLO-MIN106 --kit SQK-RNA002 --disable_pings -q 0 --read_batch_size 4000000 --reverse_sequence on --u_substitution on -x “cuda:0 cuda:1”) **(Figure S1b)**. ONT reads were mapped to the reference genome using Minimap2 version 2.17. We used parameters recommended for ONT by the Minimap2 documentation (ax splice -uf -k14 -MD).

### Preparing reference genomes and transcriptome annotations

The human and mouse reference genomes were obtained from the ENCODE portal (GRCh38 assembly for human cell types, and mm10 for mouse). All information other than the chromosome name was removed from the FASTA headers in the reference genome files. GENCODE v29 human and GENCODE vM21 mouse comprehensive GTF transcriptome annotations were downloaded from the GENCODE portal.

Since all samples were sequenced with ERCC spike-ins and SIRVs, it was necessary to augment the reference genomes and transcriptomes with these transcripts. The sequences of the SIRVs and ERCCs (Set 3) as well as the SIRV GTF annotation were downloaded from Lexogen here: https://www.lexogen.com/wp-content/uploads/2018/08/SIRV_Set3_Sequences_170612a-ZIP.zip. To create augmented reference genomes, we concatenated each of the human and mouse fasta files with SIRV_ERCCs_multi-fasta_170612a.fasta and SIRV_isoforms_multi-fasta_170612a.fasta.

Additional processing was needed before adding the SIRV and ERCC transcripts to the human and mouse annotations. Since no GTF file was provided for the ERCCs, we created one by running *merge_encode_annotations.py* on the ERCC fasta file. The SIRV isoforms (SIRV_isoforms_multi-fasta-annotation_C_170612a.gtf) were processed with the *separate_multistrand_genes.py* script so as to separate transcripts located on different strands into separate genes. Next, we ran *talon_reformat_gtf* (TALON utility) on the ERCC and SIRV GTFs in order to add in explicit gene and transcript lines needed by the TALON program. These reformatted GTF files were then concatenated to the end of the human and mouse GENCODE annotations.

### TALON pipeline

Following alignment to the genome, reference-based error correction was performed on the PacBio FLNC and ONT reads using TranscriptClean v2.0.2 (available on GitHub at https://github.com/mortazavilab/TranscriptClean). Reference splice junctions were derived from the GENCODE annotations using TranscriptClean accessory script *get_SJs_from_gtf.py*. For the human runs, we used VCF-formatted NA12878 truth-set small variants from Illumina Platinum Genomes to run TranscriptClean in variant-aware mode (--canonOnly + defaults). For the mouse datasets, we ran TranscriptClean without a VCF file (--canonOnly + defaults). By using the –canonOnly flag, we omitted any reads that still contained one or more un-annotated noncanonical splice junctions from the output.

After TranscriptClean, we ran the TALON module **talon_label_reads** on each corrected SAM file in order to compute the fraction of As following the end of each read alignment. We set the –ar parameter to 20 bp to match the length of the T sequence used in PacBio’s oligo-dT primer for poly-(A) capture. All TALON steps (including this one) were run with version 5.0. TALON and accompanying documentation are available from https://github.com/mortazavilab/TALON.

Human and mouse TALON databases were initialized from the GENCODE v29 and GENCODE vM21 + SIRVs/ ERCC annotations using the **talon_initialize_database** module from the TALON package (parameters: --l 0 --5p 500 --3p 300). To annotate the GM12878 PacBio and ONT reads, we created a configuration file with all four datasets in it and ran the **talon** module on this file along with the human TALON database (parameters: --cov 0.9 --identity 0.8). To annotate the mouse cortex and hippocampus reads, we created a configuration file with all four datasets in it and ran the **talon** module on this file along with the mouse TALON database (parameters: --cov 0.9 --identity 0.8).

To perform long read quantification, transcript abundance matrices were extracted from the TALON databases using the **talon_abundance** module. We used the unfiltered abundance files for all gene-level expression analyses (omitting genomic transcripts). To perform transcript-level analyses, we first used the **talon_filter_transcripts** utility to generate celltype and experiment-specific transcript whitelists (parameters: --maxFracA 0.5 –minCount 5). This filtering process selected for transcript models that were 1) known in GENCODE/SIRV/ERCC, or 2) reproducibly detected at least 5 times in each specified dataset. Reads with > 0.5 fraction As (as specified by **talon_label_reads**) were omitted when computing this read support. We generated separate whitelists for the PacBio GM12878, ONT GM12878, PacBio cortex, and PacBio hippocampus dataset pairs. The resulting whitelists were used to generate filtered abundance files for transcript quantification (using **talon_abundance)**, as well as custom filtered GTF annotations (using **talon_create_GTF**).

Further details and custom scripts for data visualization are available on GitHub (https://github.com/mortazavilab/TALON-paper-2020).

### PacBio vs. Illumina short read comparison

Illumina short-read RNA-seq reads from GM12878 were downloaded from the ENCODE portal in the fastq format (accession ENCSR000AEH). Quantification against the GENCODE v29 annotation was performed on each biological replicate using Kallisto^16^. The log fold changes between PacBio and Illumina counts for each GENCODE gene/transcript were computed using the exact test method in EdgeR (v3.28.1) following filtering of lowly expressed genes/transcripts and normalization. Bonferroni correction for multiple testing was performed on the resulting p-values. Genes/transcripts were considered significantly different in the two platforms if adjusted p < 0.01 and abs(log2FC) > 1.

In addition, we computed the gene-level Spearman correlations between each long-read technology and Illumina for GM12878 for genes that were detected by both platforms. To do this, we first averaged the expression (in TPM) of each gene across biological replicates by platform. For Illumina, this meant averaging the Kallisto TPM results across replicates. For PacBio and ONT, we computed the gene-level TPMs from the unfiltered TALON abundance tables for each dataset (excluding genomic transcripts and novel genes), then averaged the replicates.

### Comparison of PacBio and ONT transcriptomes

We calculated gene quantification using the unfiltered TALON abundance files with genomic transcripts removed. For transcript quantification, we included transcript models in the union of the PacBio and ONT filtering whitelists.

### CAGE analysis

Robust human CAGE peaks were downloaded from FANTOM5 in the BED format^12^. The genomic coordinates were mapped from hg19 to hg38 using the UCSC genome browser LiftOver tool^49^. We obtained the start site of each long-read transcript model from our GTF transcriptomes, then used Bedtools to ascertain whether any CAGE peak overlapped the 100 bp region immediately up or downstream of each TSS^50^.

### Computational Poly(A) motif analysis

Each GTF transcript model was converted to BED format. We extracted the DNA sequence of the last 35 bp in each transcript using the reference genome (GRCh38 assembly for human cell types, and mm10 for mouse), then searched for the presence of a known 6-mer poly(A) motif as described in Anvar *et al.*, 2018^51^.

### RNA-PET analysis

RNA-PET clusters for GM12878 were downloaded in the BED format from the ENCODE portal (accession ENCFF001TIL). The genomic coordinates were mapped from hg19 to hg38 using the UCSC genome browser LiftOver tool^49^. We obtained the start and end site of each long-read transcript model from our GTF transcriptomes, then used Bedtools to check whether any pair of RNA-PET clusters was located within 100 bp of the start and end^50^.

### Mouse Hippocampus and Cortex data analysis

Gene and transcript abundances were calculated as described above. For differential transcript expression analysis, we used EdgeR (v3.28.1) and adjusted the resulting p-values using the Bonferroni method. Transcripts with abs(log2FC) > 1 and an adjusted p-value < 0.01 were considered significantly differentially expressed. We used a custom script to identify genes that had higher novelty counts (NIC+NNC) separately for cortex and hippocampus and identified the overlapping genes. The UCSC genome browser was used to visualize transcripts colored according to their novelty.

## ACKNOWLEDGMENTS

We would like to thank Melanie Oakes at UC Irvine Genomics High-Throughput Facility (GHTF) for her help with PacBio sequencing as well as the entire ENCODE DCC for help implementing the TALON pipeline at the ENCODE portal. This work was supported in part by grants from the National Institutes of Health (UM1HG009443 to A.M. and B.W. as well as R01AG060148 and U54AG054349 to A.M and A.T.).

## Supplementary Figure Legends

**Figure S1: Platform-specific data processing performed prior to running TALON pipeline. a)** Sequencing and preprocessing of PacBio Sequel data. The Lima/Refine step in particular is important because it removes reads that did not receive a full sequencing pass and orients the remaining reads to the correct strand. **b)** Sequencing and preprocessing of ONT direct-RNA data. Since the RNA itself is sequenced poly(A) first, no additional read orientation steps are required.

**Figure S2. Performance of TALON filtering on SIRV transcripts sequenced with PacBio Sequel II. a)** Number of SIRV-aligned reads assigned to each transcript novelty category in the GM12878 Rep1 and Rep2 datasets after TALON filtering. **b)** Number of distinct transcript models called per novelty category from the SIRV-aligned reads after TALON filtering. Union of GM12878 Rep1 and Rep2 is shown. **c)** Number of SIRV-aligned reads assigned to each transcript novelty category in the GM12878 Rep1 and Rep2 datasets (no filtering). **d)** Number of distinct transcript models called per novelty category from the SIRV-aligned reads (no filtering). Union of GM12878 Rep1 and Rep2 is shown. **e)** Proportion of unfiltered SIRV reads in each novelty category that display evidence of internal priming (> 50% As in 20bp window following the alignment). Union of GM12878 Rep1 and Rep2 is shown.

**Figure S3. TALON read length distributions for PacBio GM12878 datasets. a)** Rep 1. b) Rep 2.

**Figure S4. Further characterization of gene detection in GM12878 by short reads and PacBio long reads. a)** Length of known genes by differential expression category. Gene length was computed by taking the median length of all known transcripts per gene. **b)** GC content of known genes by differential expression category. Gene GC content was computed by taking the median GC of all known transcripts per gene.

**Figure S5. Length and exon count by transcript novelty type in GM12878 PacBio. a)** Read length distributions by novelty category. **b)** Number of exons per transcript model, grouped by novelty type assignment.

**Figure S6. Epstein-Barr Virus transcriptome characterization in GM12878. a)** Gene expression levels in GM12878 from the EBV chromosome and from the human chromosomes, labelled by gene novelty. **b)** Transcript expression levels in GM12878 from the EBV chromosome and from the human chromosomes, labelled by transcript novelty. Novel transcripts have been filtered for reproducibility between GM12878 biological replicates. **c)** Visualization of TALON GTF annotations in the UCSC genome browser for EBV transcripts in GM12878.

**Figure S7. Characterization of GM12878 cell line by Oxford Nanopore direct-RNA sequencing.** TALON read length distributions for Nanopore ENCODE Tier 1 cell line datasets **a)** GM12878 Rep 1 and **b)** GM12878 Rep 2. **c)** Expression level of known transcript models and reproducible ISMs in PacBio vs. ONT for GM12878 (Pearson r = 0.48, Spearman rho = 0.08).

**Figure S8. TALON and FLAIR gene detection across sequencing platforms and samples.** Proportion of genes expressed in Illumina GM12878 RNA-seq data that are also detected by TALON, FLAIR, or both in the corresponding a) PacBio and **b)** ONT long-read datasets. Genes are divided into bins based on their Illumina expression level (TPM).

**Figure S9. Reproducibility of PacBio gene and transcript expression in mouse cortex and hippocampus. a)** Expression level of known genes in each cortex biological replicate. **b)** Expression level of known transcripts in each cortex biological replicate. **c)** Expression level of known genes in each hippocampus biological replicate. **d)** Expression level of known transcripts in each hippocampus biological replicate.

**Figure S10. TALON database schema.** Relationships between the 14 tables are indicated with grey lines, and primary keys are shown in bold.

## Supplementary Tables

| Table Name | Table description |
| --- | --- |
| **Table S1** | ENCODE/GEO IDs for generated data |
| **Table S2** | Readwise TALON annotation file for PacBio and ONT GM12878 datasets |
| **Table S3** | Readwise TALON annotation file for PacBio Cortex and Hippocampus datasets |
| **Table S4** | PB GM12878 SIRV TALON abundance table |
| **Table S5** | PB GM12878 SIRV TALON abundance (filtered) |
| **Table S6** | PacBio GM12878 TALON abundance table |
| **Table S7** | PacBio GM12878 TALON abundance table (filtered) |
| **Table S8** | PacBio GM12878 TALON GTF table (filtered) |
| **Table S9** | PacBio-Illumina differential gene expression |
| **Table S10** | PacBio-Illumina differential transcript expression |
| **Table S11** | EBV TALON abundance table |
| **Table S12** | EBV TALON abundance table (filtered) |
| **Table S13** | EBV TALON GTF (filtered) |
| **Table S14** | ONT GM12878 TALON abundance table |
| **Table S15** | ONT GM12878 TALON abundance table (filtered) |
| **Table S16** | ONT GM12878 TALON GTF table (filtered) |
| **Table S17** | PacBio-ONT GM12878 TALON abundance table |
| **Table S18** | PacBio-ONT GM12878 TALON abundance table (filtered) |
| **Table S19** | PacBio-ONT GM12878 TALON GTF table (filtered) |
| **Table S20** | Known genes enriched in ONT relative to PacBio |
| **Table S21** | Known transcripts enriched in ONT relative to PacBio |
| **Table S22** | Known gene detection by TALON and FLAIR |
| **Table S23** | Known transcript detection by TALON and FLAIR |
| **Table S24** | PacBio Mouse Brain TALON abundance table |
| **Table S25** | PacBio Mouse Brain TALON abundance table (filtered) |
| **Table S26** | PacBio Mouse Brain TALON GTF table (filtered) |
| **Table S27** | Differential transcript expression in mouse cortex and hippocampus tissue |
| **Table S28** | Genes with high degree of novelty in cortex and hippocampus |

