## supplemental methods and figures for "A technology-agnostic long-read analysis pipeline for transcriptome discovery and quantification"

### Supplementary Methods

#### *Overview of TALON database*

A key novel aspect of the TALON pipeline is its use of a database to store transcript models and abundances from multiple runs, and therefore its ability to compare new datasets to this knowledge base. The database is designed to serve two major purposes:

1) To store gene, transcript, and exon attributes of the type needed to construct a GTF transcriptome annotation (i.e. their names/IDs, genomic positions, and novelty type).
2) To track the quantity and identity of the transcripts observed in each of the datasets that have been processed so far.

The underlying philosophy of TALON filtering is that as additional datasets are sequenced and added to the database, more information becomes available to differentiate between real transcripts and artifacts. Therefore, it makes sense to apply filtering to novel transcript models in the database as a post-processing step that can be revisited at any time, rather than discarding transcripts upfront during a run. Datasets can be processed back-to-back with TALON as part of a single run but can also be added successively without the need to re-analyze the earlier data since the results are already tracked in the database.

#### *TALON database structure*

Each instance of a TALON database consists of 14 tables in total in the SQLite format **(Figure S10)**. The database is initialized from a GTF-formatted transcriptome annotation such as GENCODE, which populates its 'gene_annotations', 'transcript_annotations', and 'exon_annotations' tables with the metadata from each GTF entry. Notably, these tables permit data to be entered from more than one source, recognizing for example that it is possible for a transcript to have a different name or novelty status depending on the particular annotation version consulted.

During initialization, the locations of the genes, transcripts, and exons must also be stored. Rather than placing genomic coordinates directly in the 'gene' or' transcript' tables, we considered how the database could be extended in the future to accommodate personalized genomes for human transcriptome analysis, or genomes of different mouse strains. Individual genomic coordinates are abstracted out and represented by a vertex ('vertex' table), which can have a different location depending on the instance of the genome build in the database (as denoted in the 'location' table). Exons and introns are represented as edges connecting two vertices, which means that transcripts are paths through vertices belonging to a gene. These are stored in the 'edge' table. In the future, this graph structure could be exploited by superimposing count data onto it and examining the probability of different transcripts.

The database also contains two major tables for the explicit purpose of tracking transcripts in long read datasets. The ‘abundance’ table stores the number of times each transcript was detected in each dataset, which is highly useful for quantitative comparisons. The ‘observed’ table contains a record of every long read processed by the annotation pipeline. It tracks the read length, the transcript and gene assignment of the read, and any differences from the annotation at the 5’ and 3’ ends. The latter is important because more accurate 5’ and 3’ ends are a major goal of long read transcriptomic analysis. The ‘dataset’ table tracks associated metadata for each dataset that was initially entered by the user in the TALON configuration file.

#### *Epstein-Barr Virus transcriptome analysis*

An EBV chromosome GTF annotation was obtained from https://ebv.wistar.upenn.edu/downloadstatis/ebv.custom and refined for use. PacBio GM12878 reads that mapped to the EBV chromosome from the hg38 genome build were isolated and run through TranscriptClean using splice junctions generated from the GTF, and subsequently run through TALON. Gene and transcript TPMs were calculated using previously discussed filtering methodologies.

#### *Running FLAIR on PacBio and ONT data*

FLAIR was cloned from BrooksLabUCSC/flair on GitHub on 3/12/20 (latest commit d23a9c2ef62ede402e8b23d6231784ad910ed1af). For each of the PacBio and ONT GM12878 datasets, we ran the FLAIR align and correct steps on biological replicates separately, then combined the outputs in order to run the FLAIR collapse and quantify steps using default parameters. We removed SIRV and ERCC transcripts at this point and converted the ‘counts_matrix.tsv’ output file from FLAIR Quantify to a format resembling the TALON abundance file. From there, we compared gene and transcript detection to the TALON results and to Illumina.

#### *Long-read splice junction extraction*

Post-TALON splice junctions and GENCODE annotation splice junctions were extracted from GTF files using the get_SJs_from_gtf.py script from TranscriptClean (v2.0.2).

#### *Short-read splice junction extraction*

To obtain high-confidence splice junctions from short reads, Illumina RNA-seq reads (fastq) were mapped to the reference genome using STAR v. 2.5.2a. We used the following ENCODE-recommended parameters:

```
STAR \
--runThreadN 4 \
--genomeDir genome \
 --readFilesIn illumina_1.fastq illumina_2.fastq \
--sjdbGTFfile gencode.annotation.gtf \
--outFilterType BySJout \
--outFilterMultimapNmax 20 \
--alignSJoverhangMin 8 \
--alignSJDBoverhangMin 1 \
--outFilterMismatchNmax 999 \
--outFilterMismatchNoverLmax 0.04 \
--alignIntronMin 20 \
--alignIntronMax 1000000 \
--alignMatesGapMax 1000000 \
--outSAMattributes NH HI NM MD jM jI \
--outSAMtype SAM
```

For each splice junction, the resulting file lists genomic location, strand, intron motif, whether or not the junction is annotated in GENCODE, and the amount of read support.

To ensure splice junction reproducibility, we ran each replicate separately, and the subsequent splice junctions were merged and filtered for each cell type. Splice junctions with no supporting uniquely-mapped reads were discarded, and we required non-annotated splice junctions to have at least one uniquely-mapping read in each replicate.

#### *Splice junction support by junction novelty category*

To determine the novelty of all long-read, post-TALON GM12878 splice, we first extracted each junction from the TALON GTF. Next, we used a custom script to compare each long-read junction with the splice donors and acceptors present in the GENCODE annotation. We defined three different junction categories:

- Known junctions: The exact splice donor/acceptor combination was seen in the GENCODE annotation.
- Novel in catalog: The splice donor and acceptor are seen in the GENCODE annotation, but never together in the same junction.
- Novel not in catalog: The splice donor, splice acceptor, or both fail to be seen in the GENCODE annotation.

Once the status of each junction was determined, we computed the GM12878 short-read support for each splice junction novelty category separately by platform.

**Table S1: Accessions for submitted data**

| Platform | Cell/tissue type | Replicate | ENCODE accession | GEO accession | Raw read count | Pre-mapping read count | Reads at TALON stage |
|---|---|---|---|---|---|---|---|
| PacBio Sequel II | GM12878 | 1 | ENCLB200YVA | --- | 6,061,818 | 2,137,168 | 2,040,933 |
| PacBio Sequel II | GM12878 | 2 | ENCLB735WVC | --- | 6,692,215 | 2,538,701 | 2,445,556 |
| PacBio Sequel II | Mouse cortex | 1 | ENCLB287KUK | --- | 6,404,493 | 2,843,245 | 2,777,090 |
| PacBio Sequel | Mouse cortex | 2 | ENCLB440QNX | --- | 6,549,444 | 2,643,160 | 2,578,722 |
| PacBio Sequel | Mouse hippocampus | 1 | ENCLB722NJT | --- | 7,422,892 | 2,961,269 | 2,900,630 |
| PacBio Sequel | Mouse hippocampus | 2 | ENCLB186LWF | --- | 6,943,825 | 3,124,583 | 2,858,450 |
| ONT direct-RNA | GM12878 | 1 | --- | GSM4417547 | 2,020,127 | 2,020,127 | 1,675,608 |
| ONT direct-RNA | GM12878 | 2 | --- | GSM4417548 | 2,571,101 | 2,571,101 | 1,984,953 |
| Illumina RNA-seq | Mouse cortex | 1 | ENCLB894RIO | --- | 87,966,793 | NA | NA |
| Illumina RNA-seq | Mouse cortex | 2 | ENCLB671GZH | --- | 51,152,278 | NA | NA |
| Illumina RNA-seq | Mouse hippocampus | 1 | ENCLB591DUT | --- | 61,562,264 | NA | NA |
| Illumina RNA-seq | Mouse hippocampus | 2 | ENCLB626JBH | --- | 62,561,081 | NA | NA |

**Table S22: Detection of Illumina-expressed genes by TALON and FLAIR in PacBio GM12878**

| Illumina-expressed Genes | PacBio GM12878 | ONT GM12878 |
|---|---|---|
| **Not detected** | 13,583 | 13,788 |
| **Detected by FLAIR only** | 234 | 246 |
| **Detected by TALON only** | 2,525 | 2,381 |
| **Detected by both TALON and FLAIR** | 10,459 | 10,386 |

**Table S23: Detection of known transcripts by TALON and FLAIR in PacBio GM12878**

| GENCODE transcripts | PacBio GM12878 | ONT GM12878 |
|---|---|---|
| **Detected by FLAIR only** | 471 | 923 |
| **Detected by TALON only** | 12,741 | 11,642 |
| **Detected by both TALON and FLAIR** | 14,100 | 11,891 |

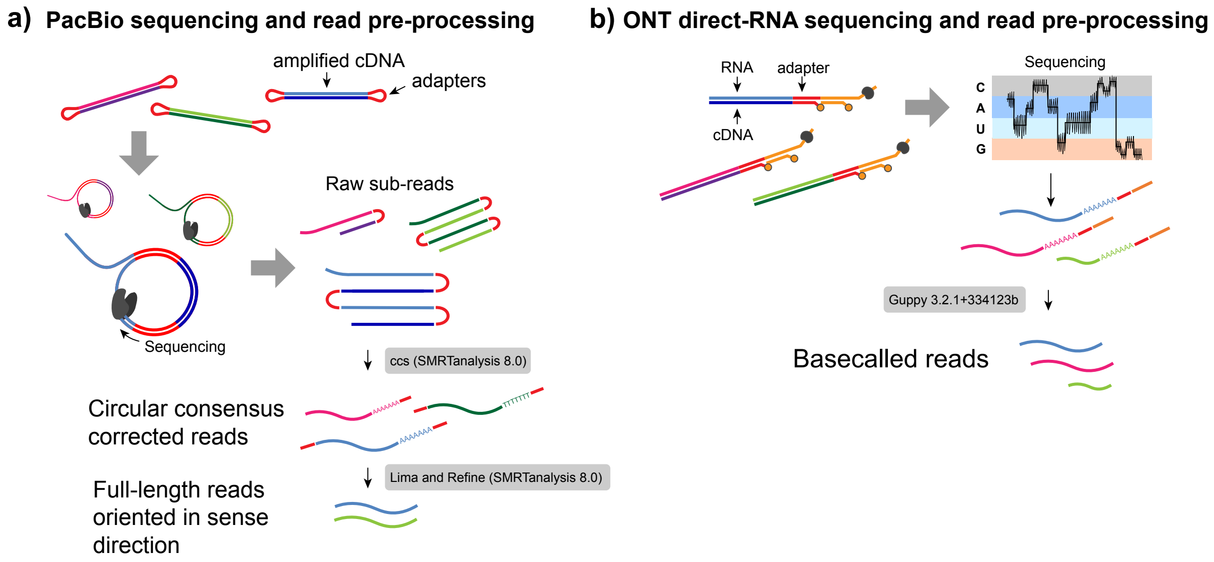


**Figure S1:** Platform-specific data processing performed prior to running TALON pipeline. **a)** Sequencing and preprocessing of PacBio Sequel data.The Lima/Refine step in particular is important because it removes reads that did not receive a full sequencing pass and orients the remaining reads to the correct strand. **b)** Sequencing and preprocessing of ONT direct-RNA data. Since the RNA itself is sequenced poly(A) first, no additional read orientation steps are required.

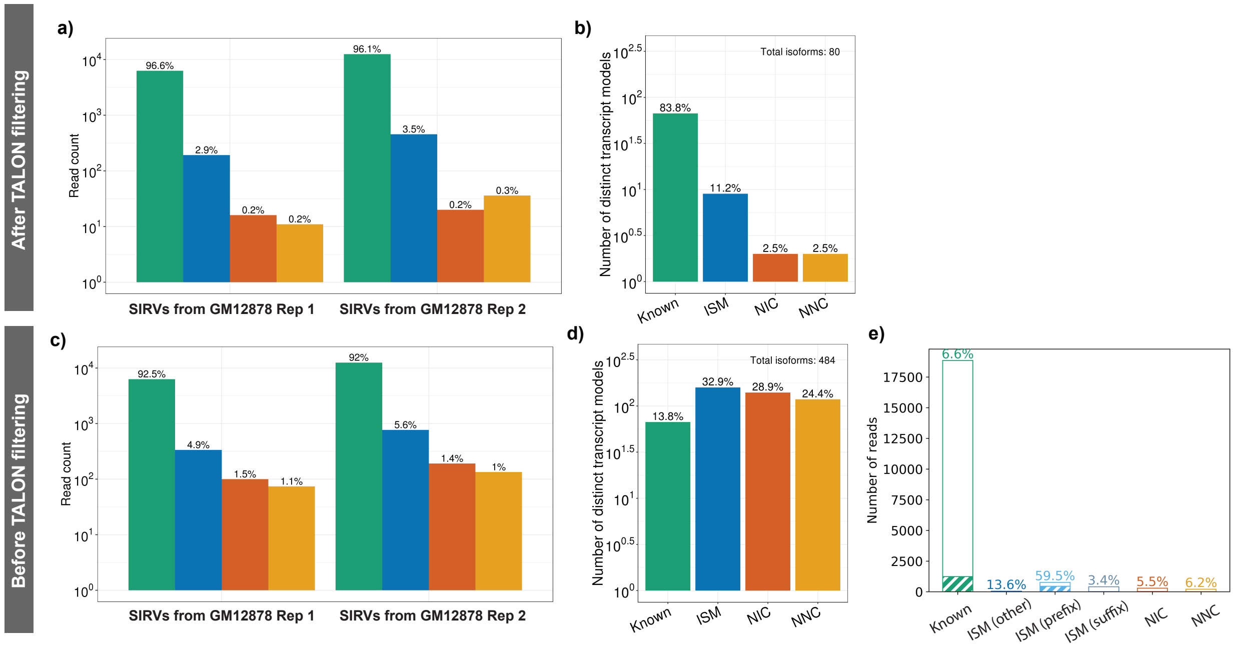


**Figure S2:** Performance of TALON filtering on SIRV transcripts sequenced with PacBio Sequel II.

**a)** Number of SIRV-aligned reads assigned to each transcript novelty category in the GM12878 Rep1 and Rep2 datasets after TALON filtering.

**b)** Number of distinct transcript models called per novelty category from the SIRV-aligned reads after TALON filtering. Union of GM12878 Rep1 and Rep2 is shown.

**c)** Number of SIRV-aligned reads assigned to each transcript novelty category in the GM12878 Rep1 and Rep2 datasets (no filtering).

Union of GM12878 Rep1 and Rep2 is shown.

**d)** Number of distinct transcript models called per novelty category from the SIRV-aligned reads (no filtering). Union of GM12878 Rep1 and Rep2 is shown.

**e)** Proportion of unfiltered SIRV reads in each novelty category that display evidence of internal priming (> 50% As in 20bp window following the alignment). Union of GM12878 Rep1 and Rep2 is shown.

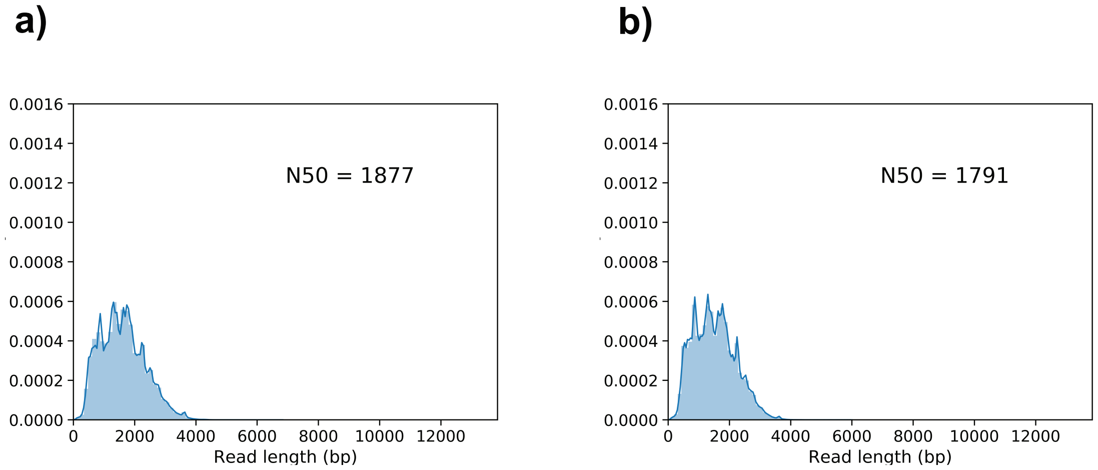


**Figure S3:** TALON read length distributions for PacBio GM12878 datasets. **a)** Rep1. **b)** Rep2.

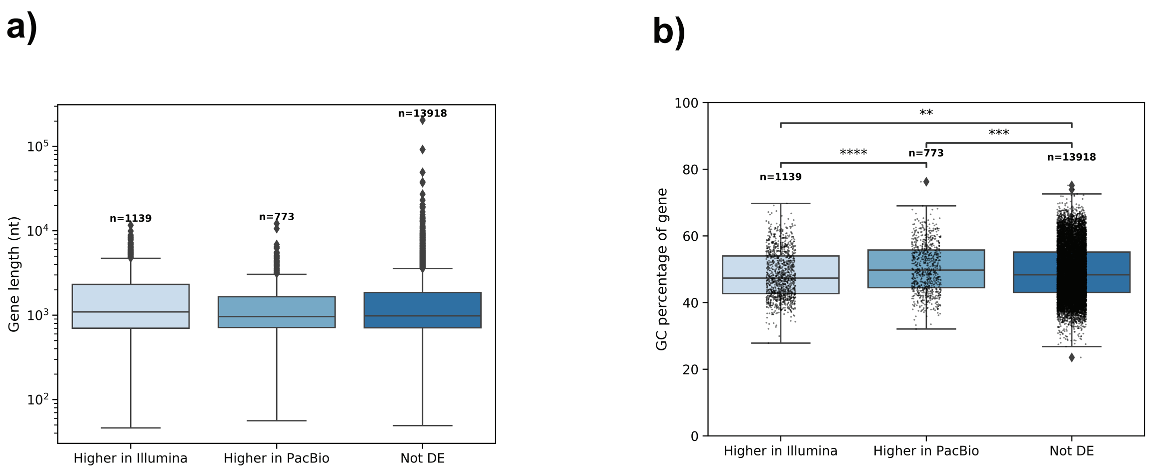


**Figure S4.** Further characterization of gene detection in GM12878 by short reads and PacBio long reads. **a)** Length of known genes by differential expression category. Gene length was computed by taking the median length of all known transcripts per gene. **b)** GC content of known genes by differential expression category. Gene GC content was computed by taking the median GC of all known transcripts per gene.

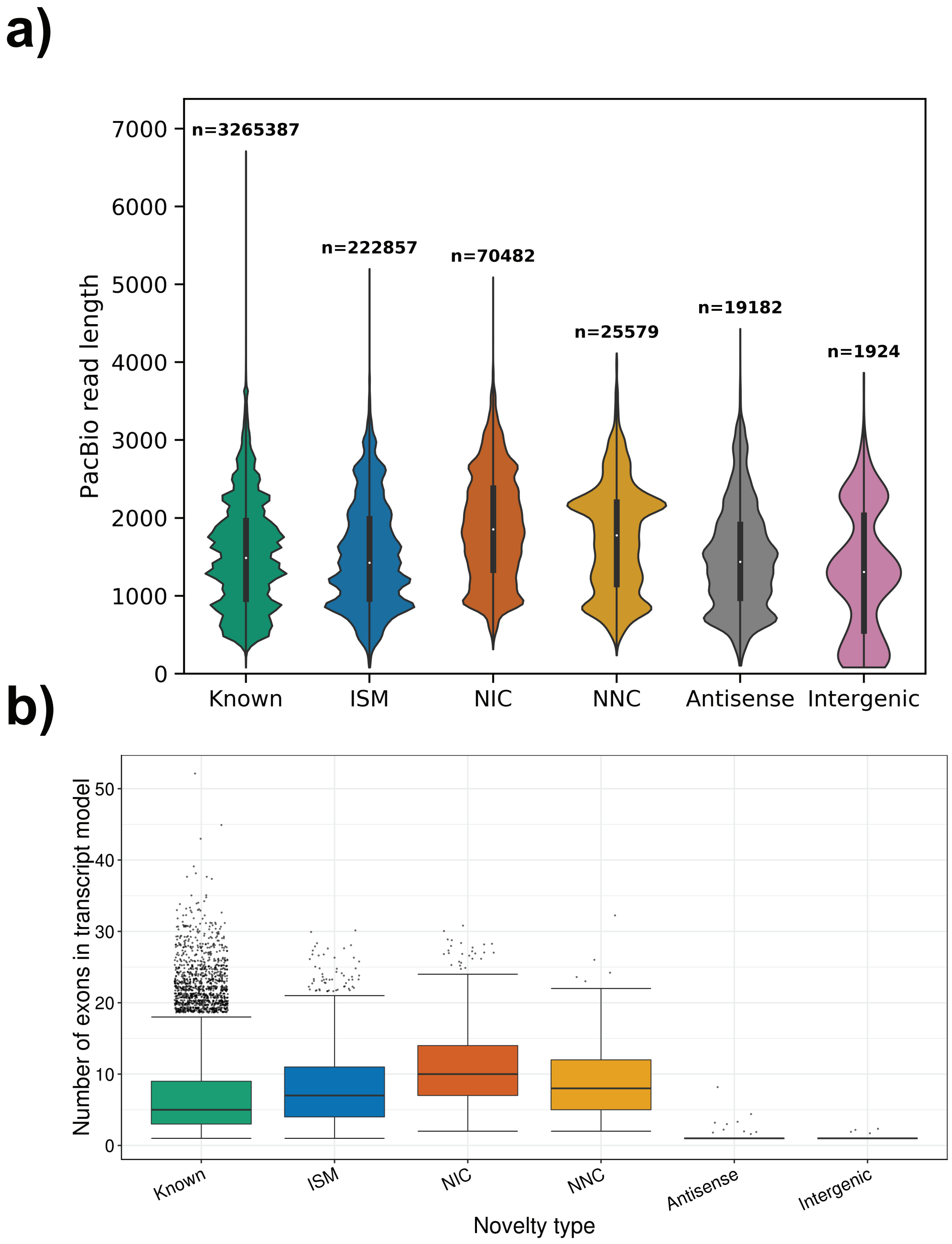


**Figure S5.** Length and exon count by transcript novelty type in GM12878 PacBio. **a)** Read length distributions by novelty category. **b)** Number of exons per transcript model, grouped by novelty type assignment.

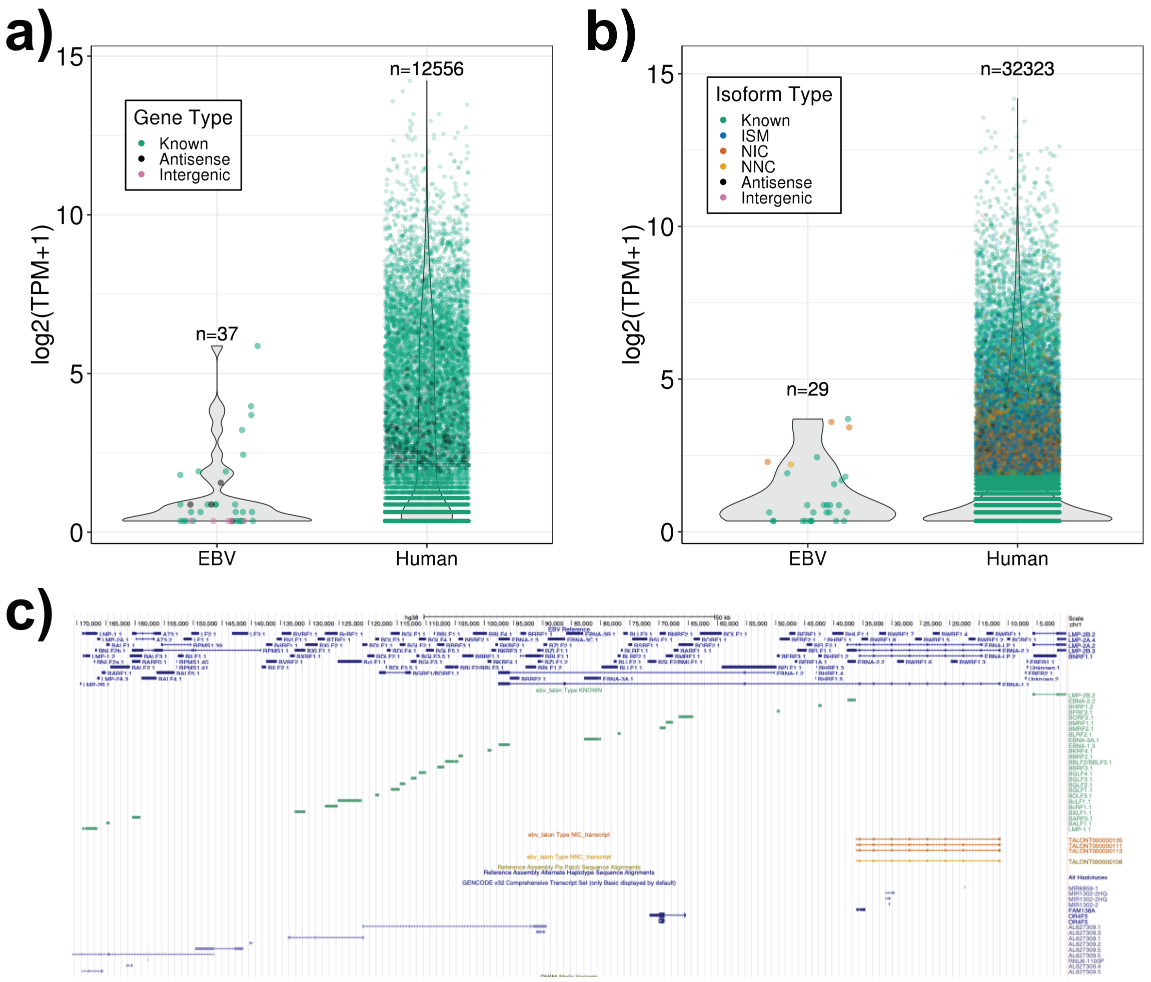


**Figure S6.** Epstein-Barr Virus transcriptome characterization in GM12878.
**a)** Gene expression levels in GM12878 from the EBV chromosome and from the human chromosomes labelled by gene novelty. **b)** Transcript expression levels in GM12878 from the EBV chromosome and from the human chromosomes, labelled by transcript novelty. Novel transcripts have been filtered for reproducibility between GM12878 biological replicates. **c)** Visualization of TALON GTF annotations in the UCSC genome browser for EBV transcripts in GM12878.

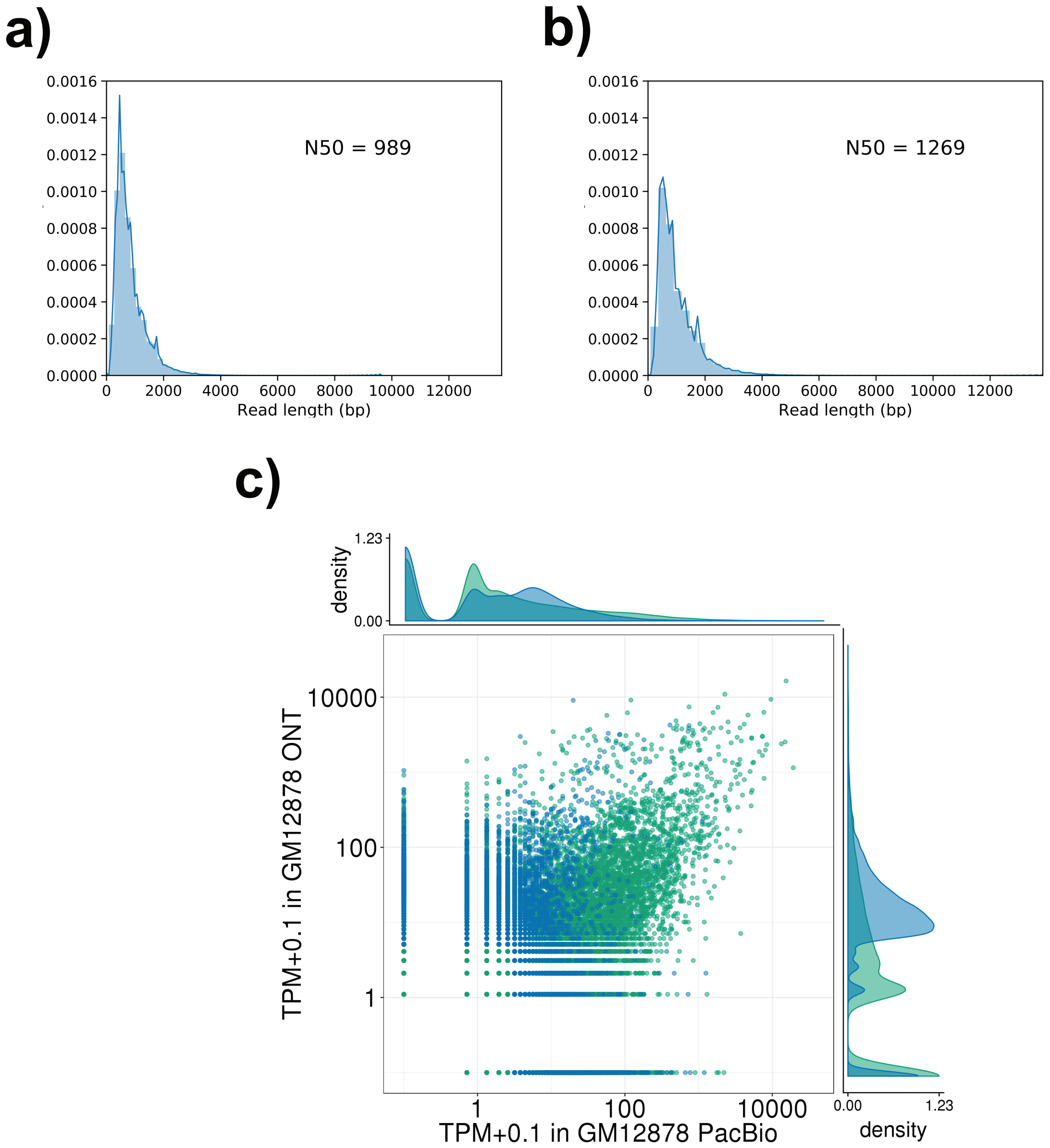


**Figure S7**. Characterization of GM12878 cell line by Oxford Nanopore direct-RNA sequencing. TALON read length distributions for Nanopore ENCODE Tier 1 cell line datasets **a)** GM12878 Rep 1 and **b)** GM12878 Rep 2. **c)** Expression level of known transcript models and reproducible ISMs in PacBio vs. ONT for GM12878 (Pearson r = 0.48, Spearman rho = 0.08).

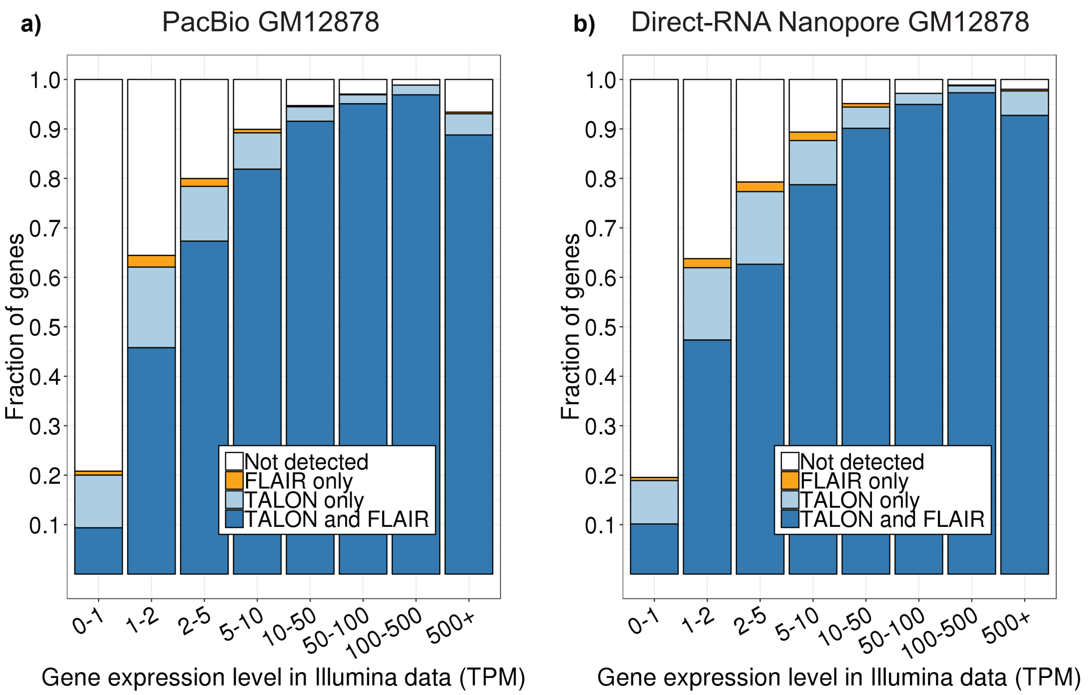


**Figure S8:** TALON and FLAIR gene detection across sequencing platforms and samples**.** Proportion of genes expressed in Illumina GM12878 RNA-seq data that are also detected by TALON, FLAIR, or both in the corresponding **a)** PacBio and **b)** ONT long-read datasets. Genes are divided into bins based on their Illumina expression level (TPM).

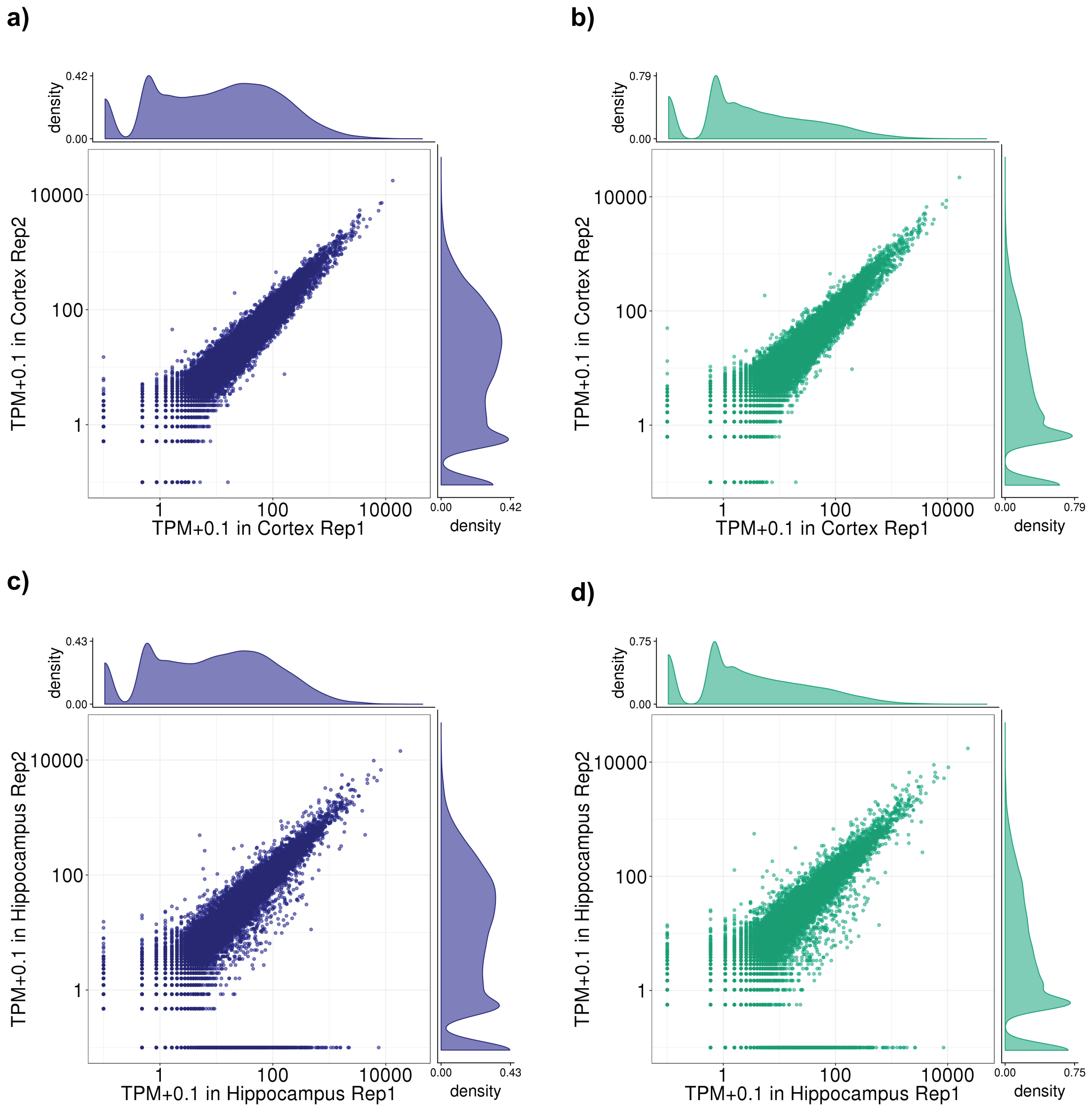


**Figure S9:** Reproducibility of PacBio gene and transcript expression in mouse cortex and hippocampus. **a)** Expression level of known genes in each cortex biological replicate. **b)** Expression level of known transcripts in each cortex biological replicate. **c)** Expression level of known genes in each hippocampus biological replicate. **d)** Expression level of known transcripts in each hippocampus biological replicate.

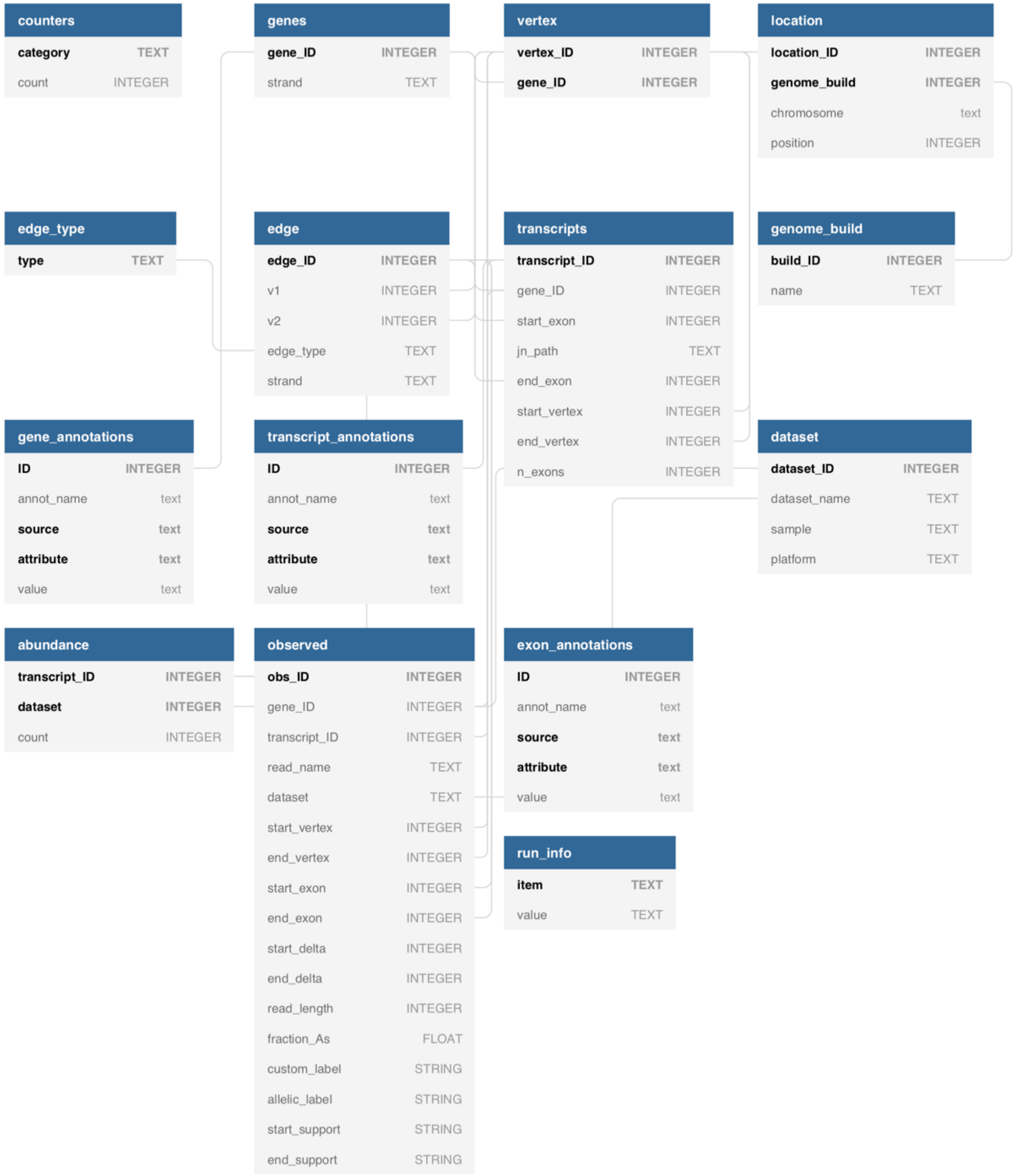


**Figure S10:** TALON database schema. Relationships between the 14 tables are indicated with grey lines, and primary keys are shown in bold.
